# Laminar CBV and BOLD response-characteristics over time and space in the human primary somatosensory cortex at 7T

**DOI:** 10.1101/2024.06.26.600746

**Authors:** Sebastian Dresbach, Omer Faruk Gulban, Till Steinbach, Judith Eck, Sriranga Kashyap, Amanda Kaas, Nikolaus Weiskopf, Rainer Goebel, Renzo Huber

**Affiliations:** Faculty of Psychology and Neuroscience, Maastricht University, Maastricht, Netherlands; Department of Neurophysics, Max Planck Institute for Human Cognitive and Brain Sciences, Leipzig, Germany; Brain innovation, Maastricht, the Netherlands; Krembil Brain Institute, University Health Network, Toronto, ON, Canada; Felix Bloch Institute for Solid State Physics, Faculty of Physics and Earth System Sciences, Leipzig University, Linńestraße 5, 04103 Leipzig, Germany; Wellcome Centre for Human Neuroimaging, Institute of Neurology, University College London, 12 Queen Square, London WC1N 3AR, UK; National Institutes of Health, Bethesda, MD, USA

## Abstract

Uncovering the cortical representation of the body has been at the core of human brain mapping for decades, with special attention given to the digits. In the last decade, advances in functional magnetic resonance imaging (fMRI) technologies have opened the possibility of noninvasively unraveling the 3rd dimension of digit representations in humans along cortical layers. In laminar fMRI it is common to combine the use of the highly sensitive blood oxygen level dependent (BOLD) contrast with cerebral blood volume sensitive measurements, like vascular space occupancy (VASO), that are more specific to the underlying neuronal populations. However, the spatial and temporal VASO response characteristics across cortical depth to passive stimulation of the digits are still unknown. Therefore, we characterized haemodynamic responses to vibrotactile stimulation of individual digit-tips across cortical depth at 0.75 mm in-plane spatial resolution using BOLD and VASO fMRI at 7T. We could identify digit-specific regions of interest (ROIs) in putative Brodmann area 3b, following the known anatomical organization. In the ROIs, the BOLD response increased towards the cortical surface due to the draining vein effect, while the VASO response was more shifted towards middle cortical layers, likely reflecting bottom-up input from the thalamus, as expected. Interestingly, we also found slightly negative BOLD and VASO responses for non-preferred digits in the ROIs, potentially indicating neuronal surround inhibition. Finally, we explored the temporal signal dynamics for BOLD and VASO as a function of distance from activation peaks resulting from stimulation of contralateral digits. With this analysis, we showed a triphasic response consisting of an initial peak and a subsequent negative deflection during stimulation, followed by a positive post-stimulus response in BOLD and to some extent in VASO. While similar responses were reported with invasive methods in animal models, here we demonstrate a potential neuronal excitation-inhibition mechanism in a center-surround architecture across layers in the human somatosensory cortex. Given that, unlike in animals, human experiments do not rely on anesthesia and can readily implement extensive behavioral testing, obtaining this effect in humans is an important step towards further uncovering the functional significance of the different aspects of the triphasic response.

## Introduction

Given the importance of our digits in daily life, unsurprisingly, a large portion of the somatosensory system is occupied by their representations. Penfield and Boldrey (1937) and Marshall et al. (1937) were the first to map the somatosensory cortex and investigate digit representations, using invasive methods in humans and macaques, respectively. Over time, the notion arose that there are 3 distinct body representations in Brodmann areas (BAs) 3b, 1, and 2 on the postcentral gyrus (for a historical overview, see Merzenich et al. (1978)). Of the 3 subregions, BA3b receives the most detailed input from the mechanoreceptors in the fingers via the thalamus and is located on the anterior bank of the postcentral gyrus. Put simply, in BA3b, the fingers are topographically organized with the thumb (D1) being found most inferolaterally, the pinky (D5) most superomedially, and the remaining digits orderly spanning the space in between (Harding-Forrester and Feldman, 2018).

Since the advent of functional magnetic resonance imaging (fMRI) and the discovery of the blood oxygen level dependent (BOLD) contrast (Ogawa et al., 1990), non-invasive investigations in humans were able to replicate the 2 dimensional maps of receptive fields in the digit-tips from the seminal papers in animals (Nelson and Chen, 2008; Sanchez-Panchuelo et al., 2010; Schweizer et al., 2008; Steinbach et al., 2022; Stringer et al., 2011; for investigations of within-digit maps, see e.g. Sanchez-Panchuelo et al. (2012), Śanchez-Panchuelo et al. (2014), and Schweisfurth et al. (2014)). Furthermore, the quantitative measurements allowed researchers to extend these findings by, for example, exploring structure-function relationships (e.g. differences in cortical thickness between functionally defined sub-regions of the primary somatosensory cortex [Śanchez-Panchuelo et al., 2014]).

In the last decade, advances in hardware (most notably ultra high field fMRI at or above 7 tesla [T]), have opened the possibility of non-invasively unraveling the 3rd dimension of digit maps cortical layers. It has been long known from the animal literature that a multitude of processes (e.g. integration of feedforward and feedback processing, predictions, attention) involve cortical layers to different degrees. Therefore, investigating laminar computations is crucial for further understanding somatosensory perception in humans (Dumoulin et al., 2018; Kuehn and Pleger, 2020). Insights into laminar digit representations in BA3b solely using BOLD measurements are sparse (for one example, see Liu et al., 2023). However, Puckett et al. (2017, 2020), Schluppeck et al. (2018), and Kalyani et al. (2023) have described the functional BOLD responses in BA3b using sub-millimetre resolution (which is sometimes regarded as a prerequisite for laminar imaging [Huber et al., 2015]). The lack of attempts using BOLD to investigate laminar processing in the somatosensory domain may, at least in part, be explained by the fact that the BOLD contrast suffers from the well known draining vein effect (Turner, 2002). Specifically, changes in blood oxygenation related to neuronal activation are strongest on the venous side of the vascular tree, which transports blood away from the site of activation. Thereby the resulting BOLD signal is displaced with respect to the active neurons (Olman et al., 2007). At high resolutions, this effect limits the effective spatial specificity of BOLD, despite small voxel sizes (De Martino et al., 2013; Polimeni et al., 2010).

To mitigate the draining vein effect in laminar fMRI, acquiring data using additional nonBOLD contrasts has been proposed, due to their usually higher specificity (for a review, see Huber et al., 2019). Vascular space occupancy (VASO; Hua et al., 2013; Lu et al., 2003) and more specifically slice-selective slab-inversion (SS-SI) VASO (Huber, 2014; henceforth referred to as VASO), is a promising cerebral blood volume (CBV) sensitive technique due to its comparatively high specificity to the microvasculature and, hence, the underlying neural activity (Huber et al., 2019). While the increase in specificity comes at the cost of sensitivity, VASO offers a measure of functional brain changes that is less dominated by large veins. Furthermore, as the VASO sequence provides both BOLD and CBV-weighted images, it is a tool for non-invasively investigating neurovascular coupling in humans (Huber et al., 2014).

To date, laminar VASO responses in the human primary somatosensory cortex have been described in several papers and conference abstracts (Dresbach et al., 2023; Yang et al., 2019; Yu et al., 2019, 2021; for VASO studies without laminar results in the somatosensory cortex, see de Oliveira et al., 2023; Huber et al., 2020, 2023a, 2023b). However, in all of these studies either fingers were stimulated in a specific order to probe prediction effects (e.g. Yu et al., 2019), did not investigate responses to stimulation of individual digits (e.g. Dresbach et al., 2023), and/or did not investigate the temporal evolution of the responses (e.g. Yang et al., 2019). As a result, the spatiotemporal VASO response across layers in the human primary somatosensory cortex to the stimulation of individual fingers is still unknown. Therefore, to investigate the laminar and temporal response-characteristics in the human primary somatosensory cortex, we acquired depth-resolved BOLD and VASO data in the somatosensory cortex using 7T (f)MRI, while passively stimulating individual finger tips.

## Methods

### Participants

Eleven healthy participants (age range: 26-46 years; mean: 31 years, 8 male, 3 female, 7 right handed) with no neurological damage participated in the study after giving written informed consent. The experimental paradigm was approved by the local Ethics Review Committee for Psychology and Neuroscience (ERCPN) at Maastricht University.

### Imaging parameters

All participants underwent scanning using a ”classical” 7T Magnetom whole body scanner (Siemens Healthineers, Erlangen, Germany), equipped with a 1-channel Tx, 32-channel Rx head coil (NOVA Medical, Wilmington, DE, USA) at Scannexus (Maastricht, The Netherlands). Functional scans were performed with a 3D-EPI (Poser et al., 2010), SS-SI VASO (Hua et al., 2013; Huber, 2014; Lu et al., 2003) sequence and a nominal spatial resolution of 0.75 × 0.75 × 1.29 mm (22 slices, inversion time [TI] = 1452 ms, effective time of repetition [pairTR] = 3850 ms, volume acquisition time [volTR] = 1601 ms, time between excitation pulses in 3D readout module [shotTR] = 67 ms, echo time [TE] = 25 ms, partial Fourier factor = 6/8 using projection onto convex sets [POCS; Nakamura et al., 2016] reconstruction with 8 iterations, FLASH GRAPPA 3 [Talagala et al., 2016], varying flip angle [FA] between 26 and 80°, read bandwidth = 1064 Hz/Px, 1st phase encoding direction along anterior-posterior axis, field of view [FOV] = 162 mm and 122 mm in read and [1st] phase encoding directions respectively). We used the vendor-provided ‘improved GRAPPA WS’ algorithm with at least 1000 fold overregularization and small GRAPPA kernels of 2 × 3 (phase x read). Slice position and orientation were chosen individually for each participant. Specifically, the FOV covered the entire postcentral gyrus without fold-over artifacts, while optimizing resolution perpendicular to its anterior bank (see Figure 1A, similar to the approach of Huber et al., 2020 in M1).

**Figure 1:**
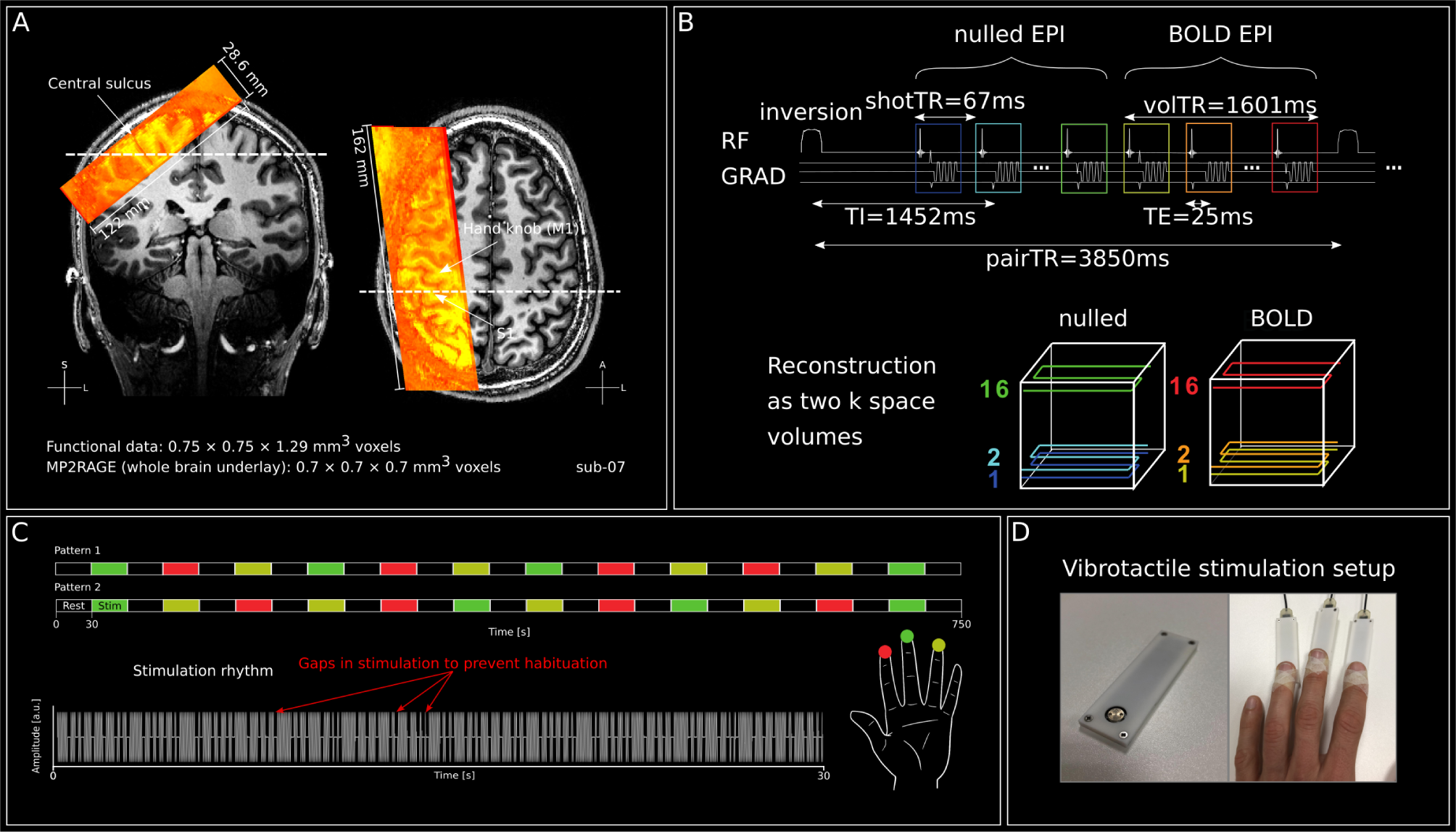
Functional data acquisition and stimulation. A. Functional EPI data (warm colors) overlaid on MP2RAGE uni image (sub-07). The fMRI slab covered the hand area of the right postcentral gyrus (right) and slices were aligned perpendicularly to the central sulcus to maximize resolution along cortical gray matter in the postcentral gyrus (left). The location of the respective other slice is indicated by the striped line in each view. **B** Functional EPI data acquisition details giving a schematic overview of relevant imaging parameters. Note: nulled and BOLD images are acquired in an interleaved fashion. Both volumes are later used to derive the ‘VASO’ images. **C** Timing of the stimulation patterns (top). For participants undergoing more than one run of stimulation, we used two stimulation patterns in an alternating fashion to decrease effects of expectancy. Furthermore, to prevent habituation over long periods of stimulation, we interspersed short breaks in the stimulation pattern (bottom). **D** Means of stimulation. Left: A single piezoelectric stimulation device. The silver disc vibrates at *∼* 25 Hz (see pattern in Figure 1C lower part) and stimulates the fingertips. Right: 3 devices are attached to the index, middle and ring fingers of the left hand with medical tape. 240 slices, TI1 = 900 ms, TI2 = 2750 ms, TR = 5000 ms, TE = 2.47 ms, FA1/FA2 = 5°/3°, bandwidth = 250 Hz/Px, GRAPPA acceleration factor = 3, FoV = 224 × 224 mm) were acquired using MP2RAGE (Marques et al., 2010). Second, 3 high-resolution anatomical images (0.5mm isotropic, 60 slices, TI1 = 900 ms, TI2 = 2750 ms, TR = 6000 ms, TE = 4.02 ms, FA1 = 6°, FA2 = 7°, bandwidth = 140 Hz/Px, acceleration factor = 2, FOV = 160 × 160 mm) were acquired with the same slab orientation as the functional data, also using MP2RAGE (Marques et al., 2010). A complete set of the scan protocols is available on: <https://github.com/sdres/sequences/tree/master/S1\_templates> and the sequences are available for VB17B-UHF via SIEMENS C2P.

Two types of anatomical images were acquired. First, whole-brain images (0.7 mm isotropic,

### Experimental Procedure

During 1-3 stimulation runs per participant (12.5 minutes each), 3 digit-tips of the left hand (index-, middle- and ring-finger, in the following referred to as D2, D3 and D4, respectively) were stimulated in a pseudorandom order (30s on-off block design, 4 repetitions/digit, Figure 1C) by means of a piezoelectric vibrotactile stimulator (Figure 1D, mini PTS system, Dancer Design, UK). The stimulation setup was controlled in Presentation® software (Version 16.0, Neurobehavioral Systems, Inc., Berkeley, CA, www.neurobs.com). A psychoPy implementation is available on: <https://github.com/layerfMRI/Phychopy\git>. Specifically, there were various stimulation patterns, with the 2 most common being displayed in Figure 1C. For participants that underwent one stimulation run, the first order was chosen, while for subjects that underwent multiple runs of stimulation, we alternated between the two protocols in order to reduce effects of expectancy. In 3 individual runs, a different pattern was used with the same number of repetitions per digit (patterns not shown). To further reduce habituation effects within a block of stimulation, we interspersed the sinusoidal stimulation pattern (25 Hz) with short interruptions (Figure 1C - bottom).

For one participant (sub-12, see Supplementary Table 1), we also stimulated the thumb (D1) and pinky (D5) of the left hand in addition to D2-D4. Increasing the duration of each run to accommodate the stimulation of additional digits would have been detrimental for participant comfort. Therefore, we reduced the run duration by only including 2 repetitions/digit in each run (resulting in a run duration of 10.5 minutes, including 30 seconds of initial rest; see Supplementary Figure S1). To account for the reduction of repetitions per digit, we acquired 4 runs of stimulation for this participant. Also here, we generated 2 stimulation patterns. For run-001 and run-002 we used pattern 1, while for run-003 and run-004 we used pattern 2. The investigation of all 5 digits is beyond the scope of the current manuscript. Therefore, we only used the statistical maps resulting from the stimulation of D2-D4.

Finally, we acquired resting state data with a run duration of either 10 (6 participants) or 20 (5 participants) minutes. Participants were instructed to lay still and keep their eyes open. The results of the resting state analysis are beyond the scope of this manuscript and will not be discussed further.

### Anatomical Data Processing

The high-resolution anatomical MP2RAGE uni images (0.5 mm isotropic) were co-registered using rigid transformations in ANTsPy (Avants et al., 2011; v0.3.3s) and subsequently averaged. Signal differences due to inhomogeneities of the radio-frequency field were corrected using *n4 bias field correction* as implemented in ANTsPy. To limit computing-demands, we reduced our anatomical data to a visually defined box around the postcentral gyrus for each participant individually. After this data-reduction, we spatially upsampled the data with a factor of 3, resulting in a nominal voxel size of *∼*0.16 mm isotropic.

To estimate gray matter borders, an initial tissue segmentation was performed using standard parameters in FSL FAST (Zhang et al., 2001; v6.0.5). Crucially, the resulting segmentations were manually improved using ITK-SNAP (Yushkevich et al., 2006; v3.8.0) by an expert (author S.D.) and quality controlled by another expert (author O.F.G.). Special care was given to the region on the postcentral gyrus opposite of the ‘hand-knob’ of the primary motor cortex. The resulting segmentation was used to estimate cortical depth levels using the *LN2 LAYERS* program as implemented in LayNii (v2.6.0; Huber et al., 2021a). Specifically, we estimated 11 layers using the equivolume principle.

### Functional Data Processing

Initially, the concomitantly acquired nulled (CBV-weighted, with BOLD contamination) and not-nulled (henceforth referred to as BOLD) time series of each run were separated and further processed individually. In order to remove timepoints in which a steady state wasn’t reached yet, we replaced the first, second and third volumes with the fourth, fifth and sixth volumes, respectively. In the following, motion correction was performed using ANTsPy (Avants et al., 2011; v0.3.3s). Specifically, time courses were corrected for within-run motion by aligning each volume to the mean of the first 3 volumes, using rigid transformations and second order spline interpolation. Thanks to the inherent T1 contrast due to the inversion recovery nature of the VASO sequence, we could derive run-wise T1w images in EPI-space (T1w-EPI) by computing the voxel-wise inverse variation coefficient of the concatenated nulled and BOLD timecourses. The T1w-EPIs of all stimulation runs were then co-registered to the T1w-EPI of the resting state run. Thereafter, we reapplied the motion correction for all stimulation runs while concatenating the transformation matrices from the within- and between-run registrations. Finally, we computed a T1w-EPI across all runs to maximize the signal-to-noise ratio (SNR) for the registration with the anatomical MP2RAGE data.

To limit computing demands, we also reduced our functional data to a visually defined box around the postcentral gyrus for each participant individually. After that, timecourses were temporally upsampled by a factor of 2 using AFNI’s *3dUpsample* (Cox, 1996, 7th order polynomial interpolation) and the first volume of the nulled timecourse was duplicated in order to temporally match the nulled and BOLD data. Finally, residual BOLD-contamination of the nulled timecourse was corrected by dividing the nulled by the BOLD timecourse. The BOLD-corrected nulled timecourses are henceforth called VASO.

We then registered the T1w-EPI (mean across runs if applicable) to the high-resolution MP2RAGE uni image. To guide the automatic registration process, we first manually aligned the images in ITK-SNAP and saved the transformation matrix. This initial transformation was then refined using non-linear SyN transformation and fifth order B-spline interpolation using *AntsRegistration*. Non-linear registration was used to account for the geometric distortions in the functional EPI data. Furthermore, we used a registration mask encompassing the area of digit representations opposite the hand-knob on the postcentral gyrus in which the optimization metric (cross correlation) should be optimized.

To estimate spatial stimulus-evoked VASO and BOLD responses, we conducted a (general linear model) GLM analysis in FSL (v6.05.2; Woolrich et al., 2001). Here, we used individual predictors for the stimulation of the 3 digits (5 digits for sub-12, see Supplementary Table 1) convolved with a standard haemodynamic response without temporal derivative (mean lag: 6 seconds, std. dev: 3 seconds), while applying a high-pass filter (cutoff = 0.01 Hz) but no additional smoothing. Furthermore, we used the output of *fsl motion outliers* from FSL and 6 motion regressors (3 rotations and translations) from the motion correction as additional confounds to minimize residual influences of head-motion (Power et al., 2014). A summary of motion exceeding the voxel size and motion outliers is shown in Supplementary Figure S2. In brief, a total of 9198 volumes (nulled and notnulled combined) was acquired across participants. Of those, 30 volumes (*<*0.33%) showed framewise displacements (FDs, Power et al., 2014) greater than our in-plane voxel resolution (0.75 mm), with only a single volume with a FD of *>*2 mm. For participants that underwent more than one stimulation run, a second level GLM analysis was run using fixed effects. The resulting statistical maps were then registered to the upsampled anatomical data using the linear and non-linear registration matrices estimated before.

To extract event related averages for individual layers, the stimulation data were registered to the high-resolution anatomical data in upsampled space using the linear and non-linear registration matrices estimated before. Because the data size in upsampled space would exceed our hardware capabilities (mostly RAM), we separated the functional timeseries into individual volumes and registered them to the anatomical data in upsampled space individually.

The analysis code is available online at: *<*https://github.com/sdres/s1Anfunco*>*

## Results

### Stimulation shows expected digit organization in putative BA3b

Figure 2A shows the mean images and temporal SNR (tSNR) maps for BOLD and VASO in a representative participant (sub-07). Crucially, the mean images are crisp and without major artifacts. Furthermore, the tSNR maps show higher values for BOLD than VASO, as expected. Nevertheless, both contrasts show a tSNR *>*10 which is adequate for fMRI at sub-millimetre resolutions (Huber, 2020). For a quantification of tSNR values in the postcentral gyrus on the group level and for all participants separately, see Supplementary Figure S2A. While there are indications of inflow effects (red arrows on the mean VASO image in Figure 2A), the affected voxels are outside of the region of interest (ROI) and therefore should not influence the results discussed below.

**Figure 2:**
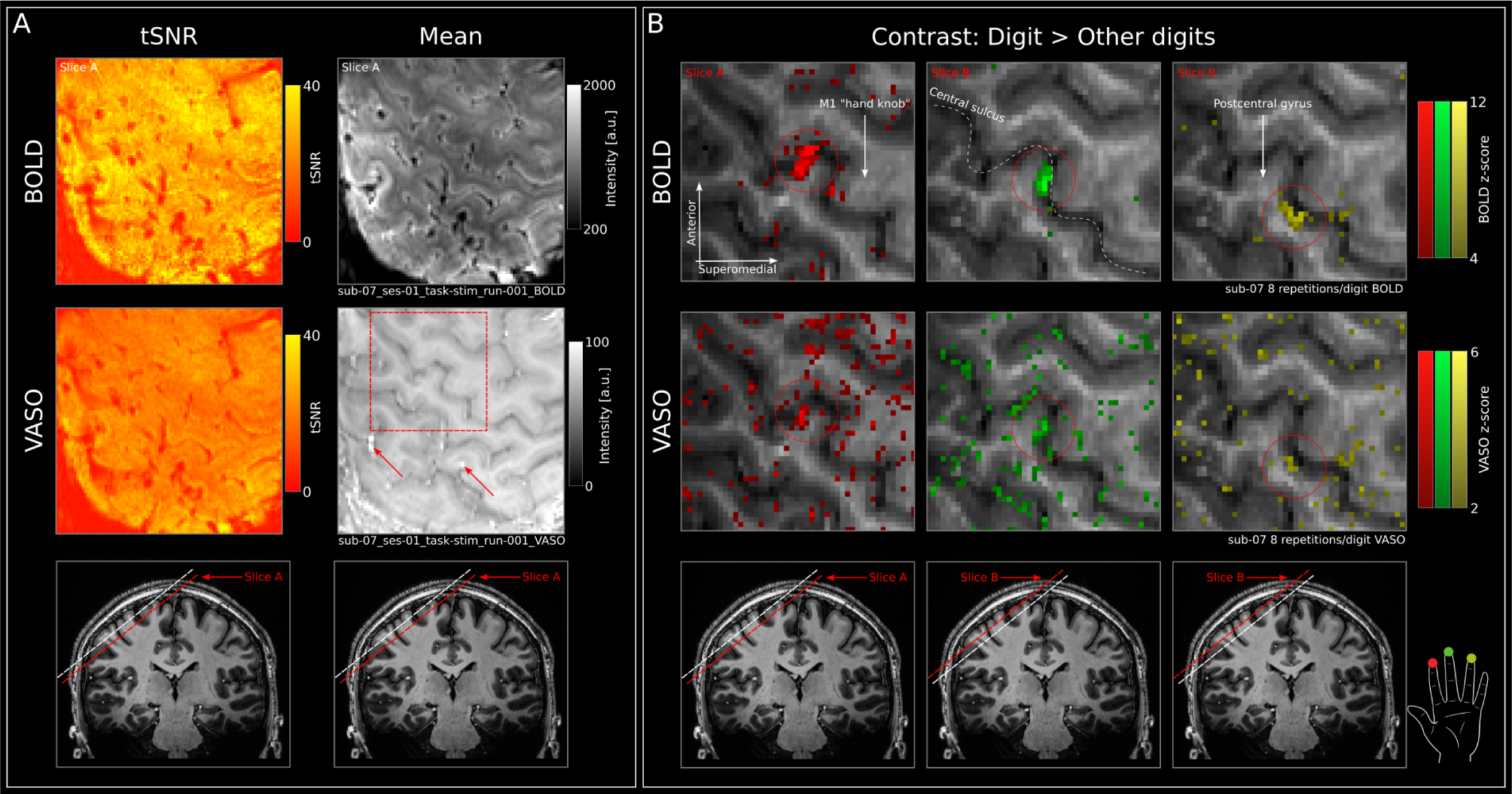
Quality assessment and stimulation results. A BOLD (upper row) and VASO (middle row) tSNR (left) and mean (right) maps of an individual run of a representative participant (sub-07). Red square in the middle right panel indicates the zoomed-in section in B, while the slice position with respect to the entire brain is indicated in the lowest row. In the ROI, we observe tSNR values >10 for both BOLD and VASO, which is adequate for analyses at sub-millimetre resolution. Mean BOLD and VASO images are crisp and show no major artifacts. Bright voxels in the mean VASO image may indicate inflow of non-inverted blood (red arrows) but are outside the ROI and can therefore be ignored. B BOLD (upper row) and VASO (middle row) stimulation results showing the statistical z-map resulting from a GLM analysis in which we contrasted each finger against all other fingers overlaid on a T1w image directly in EPI space (therefore without alignment issues). Note that we display only individual contrasts within a given map for better visual clarity. In addition, note that we did not apply thresholds based on cluster-size, to give a better feeling for the noise characteristics in the data.

Tactile stimulation of individual digit-tips (D2-D4) of the left hand evoked strong BOLD responses in the right primary somatosensory cortex of all participants (GLM contrast: respective digit *>* all other digits; e.g. for D2 in BOLD [D2: 1, D3: -0.5, D4: -0.5]). The BOLD results of one representative participant (sub-07) are shown in Figure 2B (upper row), whereas the results from all participants are shown in Supplementary Figures S3-S5. As expected, we observed stimulation induced activation in the putative BA3b on the anterior bank of the postcentral gyrus, following the known topographical organization in 10 out of 11 participants.

Specifically, we found D2-D4 to be neatly represented in order from lateral to medial and inferior to superior locations. Note that the data were acquired with an oblique slab orientation in all participants. Therefore, even when multiple finger representations are seen in the same slice, more medial representations are also more superior.

In the deviating participant (sub-10), we found two clusters for D3. One cluster is located laterally and inferior to D2. The other cluster is located between D2 and D4, as would be expected. Importantly, the “irregular” cluster was found to be larger and showed stronger activation within gray matter. Furthermore, similarly irregular patterns of digit representations have been observed in this participant with fMRI before (e.g. Steinbach et al., OHBM 2023). Therefore, we considered the “irregular” cluster to be the representation of choice in further analyses.

In the VASO data, clearly identifiable responses to the stimulation of all digits can be seen in 7 out of 11 participants (sub-06, sub-07, sub-09, sub-12, sub-14, sub-15, sub-18). Figure 2B (middle row) shows the VASO results of one participant (sub-07; for the data of other individual participants, see Supplementary Figures S3-S5). In a further 2 participants (sub-16, sub-17), responses were clearly visible for 2 out of 3 digit representations. For the remaining digit representation, clusters with increased activation in response to stimulation could be identified when taking the BOLD-activation as guidance. In the two remaining participants (sub-05 and sub-10), one digit representations could be identified in VASO when taking the BOLD activation into account. The other two representations could not clearly be picked up from the noise in the data.

Note, that for 4 participants (sub-05, sub-06, sub-09, sub-10), we only acquired one run of stimulation data, yielding 4 blocks of stimulation (30 s, each) per digit. Compared to other VASO experiments with a similar setup, this is an exceptionally small amount of data. For example, Huber et al., 2021b used 20 blocks of stimulation (30 s, each), which is almost twice as much as for the participants that underwent the most scanning in our experiment (12 blocks per digit). Therefore, absent or low activation is expected and sub-05 and sub-10 were excluded from further VASO group analyses.

### Stimulation shows expected activation across cortical depth

To quantify activation across cortical depths, we semi-automatically generated digit-specific ROIs. Figure 3A shows the procedure and resulting ROIs in 3D for a representative participant (sub-07). First, the functional data were registered to the upsampled high-resolution MP2RAGE UNI images. Figure 3A (leftmost column) shows the gray matter segmentation overlaid on the UNI image and the registered T1-EPI. It can be seen that the segmentation accurately follows the gray matter ribbon in both images, indicating the high registration accuracy. The ROIs were then generated using the following 5-step procedure:

1. From the segmentation, we estimated cortical depth levels using *LN2 LAYERS* with the equivolume option as implemented in LayNii (v2.6.0).
2. We then generated a participant-specific geodesic gray matter disc using *LN2 MULTILATER* to limit the ROI generation to the digit region on the anterior bank of the postcentral gyrus. The central point and radius of the disc was chosen individually for each participant to encompass the BOLD activation (contrast digit *>* other digits). For 10 out of 11 participants, the radius of the disc was set to 11 mm. For one participant (sub06), the digit clusters were further spread out and a radius of 12 mm was necessary (see Supplementary Figure S6).
3. The BOLD activation of each digit was thresholded at 1/3 of its respective maximum zvalue, binarized and limited to cortical gray matter by intersecting the binary digit mask with the gray matter disc (generated in step 2).
4. We used the cortical parametrization from the *LN2 MULTILATERATE* program and the binary digit masks to propagate the largest cluster of the binary digit mask across cortical depth using LN2 UVD FILTER with the -max option (Pizzuti et al., 2023). Briefly, the algorithm iteratively places a cylinder (chosen radius: 0.45 mm) around each voxel within the gray matter disc perpendicular to the cortex and gives all voxels within the cylinder the maximum value found within it. Given that we used the binarized digit map as input, voxels within cylinders containing at least one “active” voxel are labeled 1, whereas voxels within other cylinders are labeled 0. With this procedure, we assured that a similar number of voxels was included across depth-levels.
5. Finally, we removed any voxels from the digit masks that were part of multiple digit ROIs. The average extent of the resulting ROIs was 58.54, 58.76, 44.38 mm^3^ across participants for D2, D3 and D4, respectively (For digit-specific ROI volume of individual participants, see Supplementary Figure S6K).

**Figure 3:**
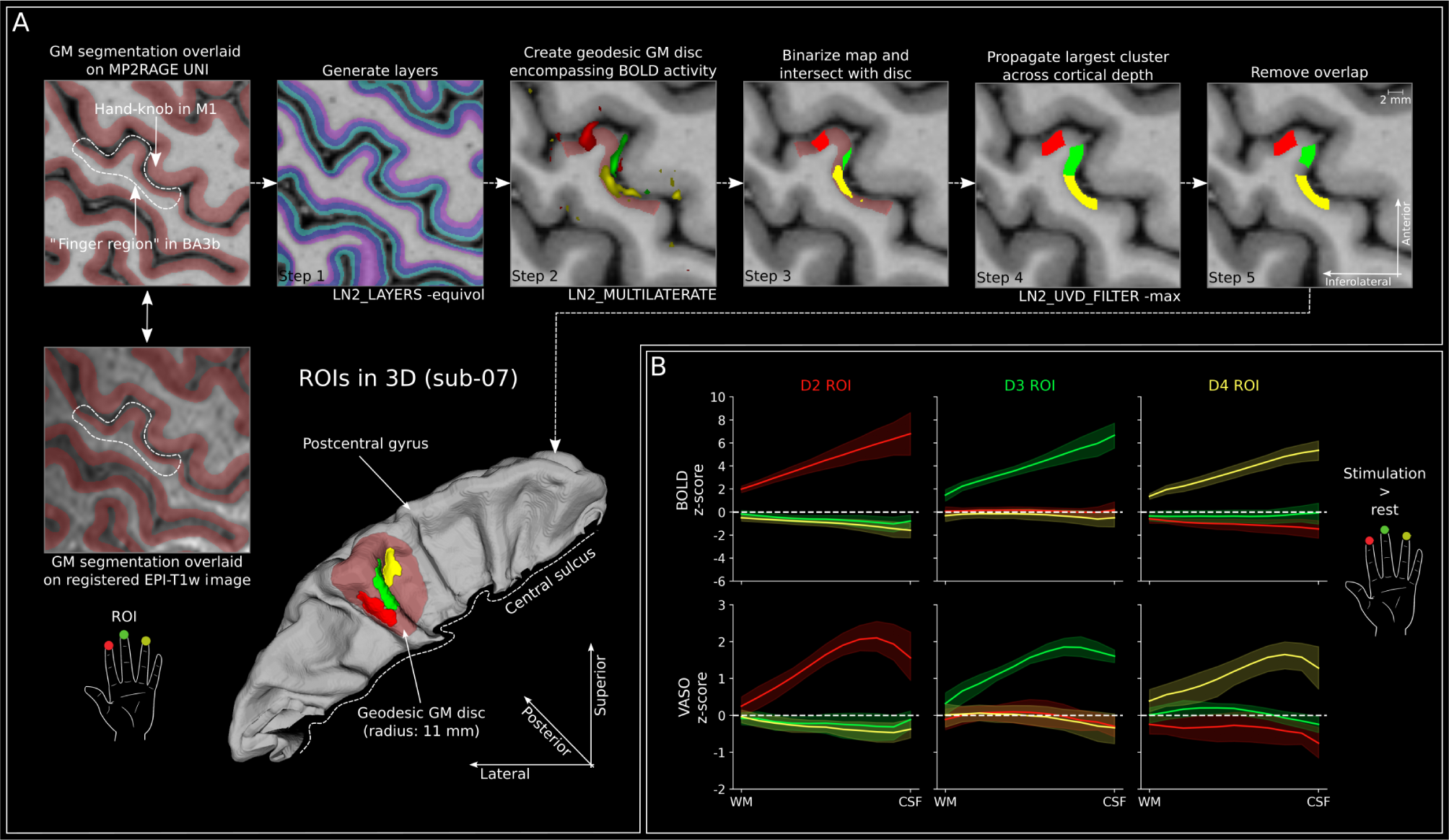
ROI generation and activation across cortical depth. A Leftmost column shows the gray matter segmentation overlaid on the upsampled MP2RAGE UNI image (nominal resolution: 0.16 × 0.16 × 0.16 mm^3^ isotropic; upper row) and the same segmentation overlaid on the registered EPI-T1w image (lower). Note that the segmentation matches the gray matter ribbon in both panels, especially in the area of interest, highlighting proper registration. Dashed lines indicate the approximate region in which the 3 fingers of interest are represented in this slice. The remaining tiles show the steps in the automatic process of generating digit-specific ROIs based on BOLD activation (GLM contrast: digit *>* other digits: e.g. for BOLD D2: [D2: +1; D3: -0.5; D4: -0.5]). Note that in step 1, we display 3 layers for simplicity. The actual number of layers from which data is extracted will be indicated for each analysis. Also note that in step 2, the statistical threshold was set to 1/3 of maximum of the respective digit-specific map. For example, if the maximum z-value across voxels for the contrast D2 *>* other digits (red color in all plots) was 12, the threshold for ROI generation was set to4. This automatic process resulted in ROIs spanning multiple slices in all participants (shown for one representative participant [sub-07] in 3D). B Stimulation-induced activity (z-scores) across cortical depth for BOLD (upper row, n = 11) and VASO (lower row, n = 9) for each digit ROI separately (GLM contrast: digit *>* rest). Values were extracted from participant-specific ROIs (example depicted for sub-07 in A using 11 depth-levels. Z-scores across cortical depth clearly show strongest BOLD activity towards the cortical surface and response peaks between middle and superficial cortical layers for VASO in response to stimulation of the preferred digit within each ROI. White dashed line indicates 0 line and shaded area indicates 95% confidence interval across participants.

The BOLD and VASO responses across cortical depth within the digit-ROIs are shown in Figure 3B (for individual participants, see Supplementary Figures S7 & S8). For each ROI, z-scores resulting from finger stimulation of each finger were extracted using 11 depth levels and averaged across participants (GLM contrast digit *>* rest). BOLD activity clearly shows an increase in activation towards the cortical surface for the preferred digit within each ROI, as expected based on the draining vein effect. VASO activity shows peaks more towards middle cortical layers and a decrease towards the surface, as expected due to the higher specificity towards underlying neuronal populations of VASO. Here, the activation closer to middle cortical layers likely reflects feedforward thalamic input arriving in cortical layer 4. Note that the ROIs in Figure 3A were based on the BOLD activation, which has a larger extent than the VASO activation due to its higher sensitivity (also reflected in higher z-scores in Figure 2B & Figure 3B). As a result, the ROIs include many voxels without substantial VASO activity and likely underestimate the sensitivity and specificity of the extracted VASO activation. Furthermore, we find slightly reduced negative responses for other digits within the D2 and D4 ROIs (activity below dashed lines in Figure 3B). Interestingly, activity for digit representations that are further away are more negative. This may hint towards surround inhibition of more distant digit representations, whereas digits that are closer are often used together and therefore not inhibited as strongly. In line with this, there is only very weak negative activity within the D3 ROI occurring with stimulation of D2 or D4, given its position centered between the two. Finally, the negative response seems to be strongest in superficial layers for both BOLD and VASO. However, this effect is rather small and may be circular due to our ROI definition based on the contrast digit *>* other digits. Therefore, it should be interpreted with caution.

### Temporal analysis

#### Event-related averages show a triphasic response to distant, non-preferred digits

In order to analyze the temporal evolution of activation across cortical depths, we extracted event related averages from 3 cortical depth levels and 3 digit-ROIs for BOLD and VASO independently. To do so, we first used the program *signal.clean* implemented in nilearn (v0.10.3; Abraham et al., 2014) to linearly detrend the voxel-time courses, regress out the motion traces and motion outliers (see section “Functional Data processing” in the “Methods” section) and compute signal changes (%) with respect to the mean of the original time series. The 3 layer-bins were generated by collapsing across layers 1-4 (deep), 5-7 (middle), and 8-11 (superficial) given by *LN2 LAYERS* to extract cortical depth profiles (see Figure 3). We chose to include the outermost layers into the deep and superficial bins because of their limited voxel counts due to the known gyral/sulcal bias in equivolume layering (i.e. superficial layers are thinner in gyri and deep layers are thinner in sulci [Bok, 1929; Waehnert et al., 2014]). The corresponding group results for BOLD and VASO are shown in Figure 4A and Figure 4B, respectively.

**Figure 4:**
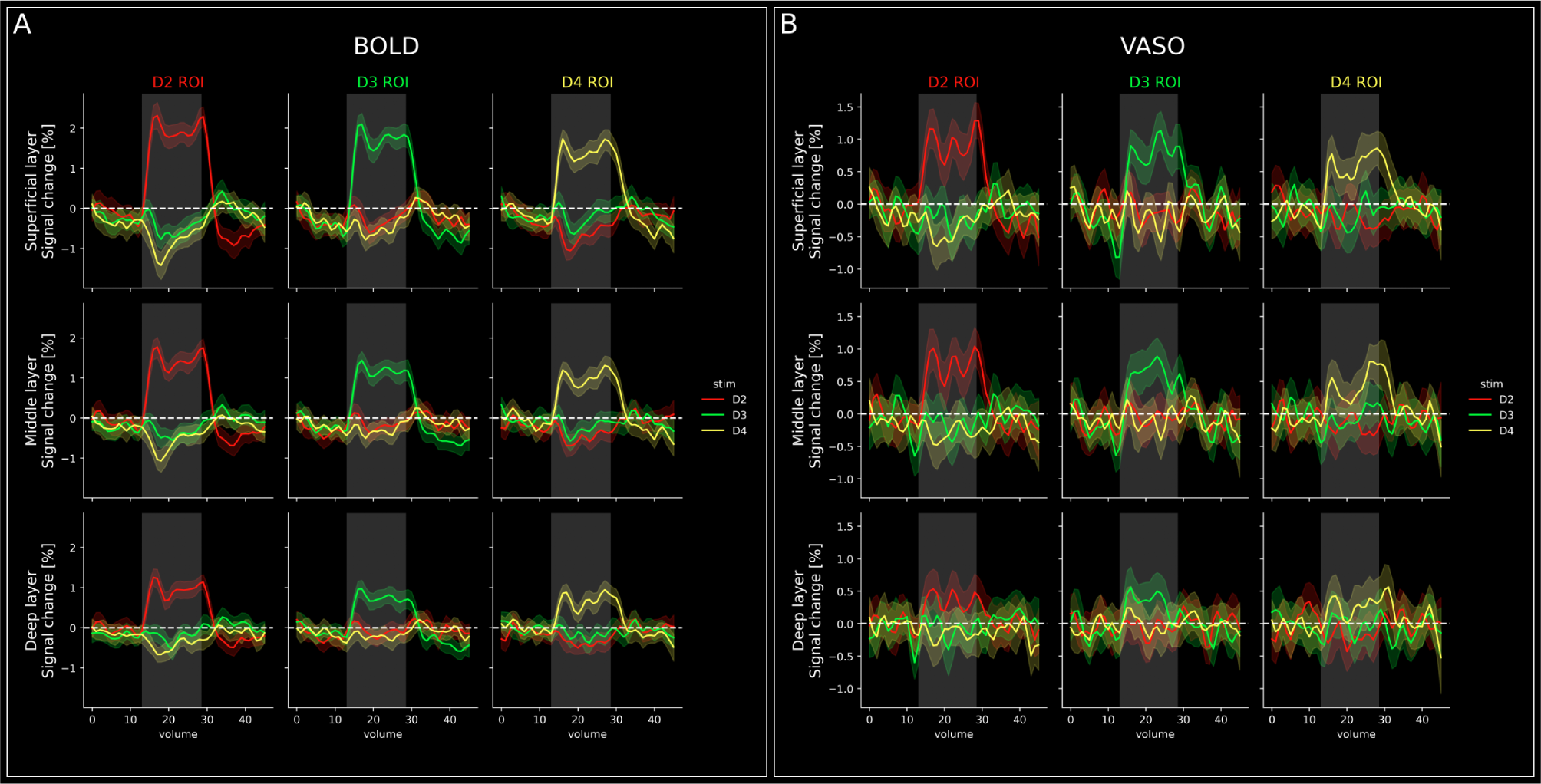
Event-related averages across cortical depths. A Event-related averages for BOLD data (n = 11) in superficial (top) middle (middle) and deep (bottom) layer compartments. Gray shaded area indicates the stimulation period. The striped line indicates the baseline, which was calculated as the voxel-wise mean of the entire run. The shaded area around lines signifies the 95% confidence interval across trials. There are 3main observations: First, as expected based on the ROI definition, strongest responses in each ROI are always found for stimulation of the preferred digit. Second, response strength to stimulation of the preferred digit increases towards the cortical surface. Third, there is a negative response to non-preferred digits in all ROIs, which seems to be stronger (more negative) for stimulation of more distant digits. B Same as A, but for VASO data (n = 9). Similar patterns can be observed, however with higher noise. Furthermore, the response is slightly less biased towards the cortical surface for preferred digits in all ROIs.

As expected based on our ROI definition, responses to stimulation of each digit are highest in the corresponding ROI for both BOLD and VASO. Furthermore, BOLD responses to stimulation of non-preferred digits seem to be slightly negative in all ROIs and across cortical depths. This is not surprising, given that we selected the ROIs based on the GLM contrast of each digit *>* other digits. Also unsurprisingly, we see highest BOLD responses to stimulation of the preferred digit in the superficial layer and the lowest responses in the deep layer of the respective ROI. VASO response amplitudes to stimulation of the preferred digits are comparable in superficial and middle layers with slightly higher responses in the superficial layer.

Smallest responses are seen in the deep layer. For the VASO z-scores across cortical depth, we observed a slight signal decrease towards cortical surface (Figure 3). The fact that we do not observe a similar effect in the event-related averages can be explained by partial voluming due to fewer layers. Furthermore, z-scoring will reduce the signal of superficial layers due to the higher noise level, which is not the case in the raw % signal change. Additionally, it could be noted that different to some previous layer-fMRI VASO studies, the gray matter segmentation (shown in Figure 3A) is performed to only include voxels that are solidly filled with gray matter only. Thus, the profiles shown here do not include signal and partial voluming of pial voxels, where BOLD and VASO are expected to diverge more strongly. Also, the decreased activation in response to stimulation of non-preferred digits is less apparent in VASO than in BOLD. However, we can see a trend in a similar direction despite the higher noise level in the VASO compared to BOLD data. Finally, we find indications of a triphasic response to non-preferred digits in BOLD. Specifically, we find a small initial BOLD signal increase, followed by stronger negative signal changes and a small positive response after stimulus cessation. The negativity is most visible in ROIs corresponding to D2 and D4 and more so in superficial, compared to deeper layers. While the initial increase does not necessarily reach positive signal changes, it constitutes a consistent increase with respect to previous timepoints.

Note that the inter stimulus interval in our experiment was 30 seconds. Therefore, the volumes 0-15 and volumes 30-45 in Figure 4 are actually overlapping. In volumes 30-45, we can see a signal decrease for non-preferred digits and remnants of an undershoot in preferred digits. Therefore, the fact that the initial response to non-preferred digits does not exceed 0 is likely due to a carryover effect from preceding trials. However, while the data comes from the same timepoints, they are still independent because the same digit was never stimulated multiple times in a row.

### The triphasic response changes as a function of distance from the activation peak

To further investigate the triphasic response of non-preferred digits, we first identified the peak BOLD voxel in response to the stimulation of each digit individually. We then defined 7 euclidean distance bins within gray matter ranging from 0-2 to 12-14 mm (in 2 mm increments) from the peak BOLD activation for D2, D3 and D4 independently (Figure 5A) using *LN2 GEODISTANCE* implemented in LayNii (v2.6.0). The area in which we defined the distance bins was limited to the geodesic disc used for the creation of ROIs (red shaded disk in Figure 3A) because the tissue segmentation was performed especially carefully here.

**Figure 5:**
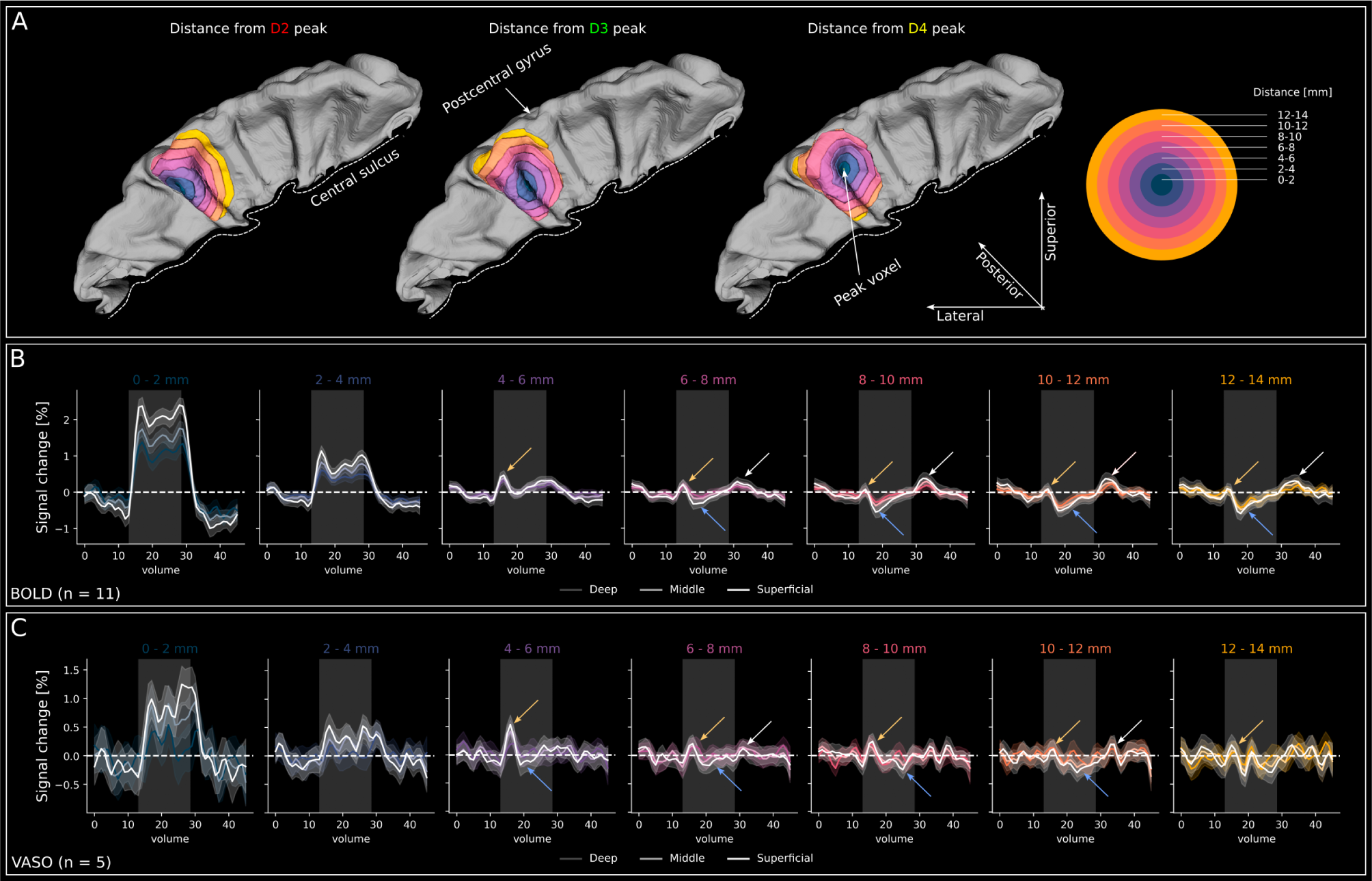
Triphasic response as a function of distance from peak activation. A 3D representations of the postcentral gyrus with euclidean distance bins from the BOLD peakvoxel of D2, D3 and D4 response (from left to right). The participant (sub-07) is the same as in Figure 2 and Figure 3). The color code of distance-rings is explained on the right. B Group-level (n = 11) BOLD event-related averages for 7 distance bins with respect to the peak voxel, collapsed over all 3 stimulated digits. The data are plotted for superficial, middle and deep layers per distance bin. Superficial layers are white in all panels, whereas middle and deep layers are color coded with increasingly darker colors of the respective distance bin (Figure 5A). The yellow arrows indicate the small initial signal increase upon stimulation. The cyan arrows indicate the following signal decrease. The white arrows indicate post-stimulus signal increase in more distant bins. Gray shaded area illustrates the stimulation period. Shaded area around lines signifies the 95% confidence interval across trials. C Same as B, but for VASO (n = 5). Note that we only plotted VASO data from participants that showed the triphasic response in the BOLD data on an individual participant level.

We then plotted group-level (n = 11) BOLD signal changes in superficial, middle and deep layers for each of the 7 distance bins independently (Figure 5B). For the 0-2 mm bin, the response shape resembles the combined responses from the digit ROIs (compare to Figure 4A). Similarly, for the 2-4 mm bin, we see activation resembling the expected response pattern from regions responding to the stimulation of preferred digits, albeit weaker. Interestingly, the 4-6 mm distance bin starts to show a small initial response (yellow arrows in Figure 5B), followed by a slight signal decrease (cyan arrows in Figure 5B). In the remaining distance bins, the initial BOLD response seems to become increasingly weaker, whereas the following negative signal changes seem to become increasingly negative. Crucially, unlike in Figure 4, the voxel selection in this analysis is almost completely independent from a GLM contrast (except for the definition of the peak voxel) but on anatomical distance from a peak voxel. Therefore, the effects cannot be explained by a voxel-selection based on response-characteristics. Finally, the more distant distance bins (*>* 6-8 mm) show a notable positive post-stimulus peak starting at the stimulus offset (white arrow in Figure 5B).

To investigate the consistency of these results, we also examined the data of individual participants (Supplementary Figures S9 and S10). We observed a pattern that resembles the group results in 5 out of 11 participants. Crucially, this seems to be related to the number repetitions of stimulation we acquired for each participant. In 5 out of the 7 participants for which we acquired *>*8 repetitions per digit, we observed a similar pattern as the group results. For 4 of the remaining 6 participants, we only acquired 4 repetitions per digit and the data quality may not be sufficient to detect these signal changes with respect to the noise in the data.

To make sure that the triphasic response we observed is not restricted to the ROIs, we performed a control analysis in which we excluded data from the digit ROIs (Supplementary Figure S11A). The results can be seen in Supplementary Figure S11B. Crucially, the triphasic response in more distant bins remains intact even if the digit ROIs are excluded. Furthermore, we observe a reduced but still substantial response in the closest distance bins. At first glance, this may seem counterintuitive given that the peak voxel lies within the digit ROIs. However, there are three major points that can explain this. First, the peak activation voxel may be close to the border of an ROI. Therefore, the 0-2 mm radius may contain non-ROI voxels.

Second, we defined the digit ROIs based on the contrast digit *>* other digits, while in the current analysis, we are estimating signal changes with respect to baseline. Signal changes with respect to baseline are usually more extensive than comparative contrasts. Third, the ROIs were defined using relatively high thresholds (z-score above 1/3 of the maximum), while the results of the current analysis do not depend on a statistical threshold. For these reasons, it is conceivable that regions outside of our ROIs show positive responses to stimulation.

To investigate whether the triphasic response is driven by the stimulation of a specific digit, we plotted the BOLD responses of one participant with high data-quality (sub-16) for individual digits separately (Supplementary Figure S11C). This means that the data going into the analysis is reduced by a factor of three. Nevertheless, indications of a triphasic response are preserved in all individual digits in this participant. Even though this is not proof that all digits contribute equally, it is evidence that at least all digits seem to contribute to some degree.

We further investigated the VASO results for the participants in which we could identify the triphasic response for more distant bins in the BOLD data. The group level VASO results (n=5) are shown in Figure 5C and the individual participants are shown in Supplementary Figure S12. On the group level, a similar triphasic response as in the BOLD data can be observed, however, it is less clear due to higher noise levels and variability across participants in the data.

### Tentative quantification of the triphasic response

We quantified the times to peak (TTP) and the respective signal changes (%) of the initial peak, trough and post-stimulus peak of the triphasic response based on the mean response across participants for distance bins *>* 4-6 mm and each layer individually (for BOLD n = 11; for VASO n = 5). The peak was defined as the highest deflection within the stimulation period, while the trough was defined as the lowest deflection after the peak and before the end of the stimulation period. The post-stimulus peak was defined as the highest deflection in a window of 8 volumes (*∼* 15.4 seconds) after stimulus offset. For the post-stimulus peak, we only took distances *>* 6-8 mm into account. This is done because we did not observe a corresponding response in the group results for smaller distances (Figure 5B). Because of the high variability across participants, we extracted information on BOLD and VASO TTPs and respective signal changes based on the mean response across participants. Therefore, we did not test the statistical significance of any laminar or distance related differences of TTPs or peak and trough magnitudes. Furthermore, as noted before, the inter-trial interval was not sufficiently long to let the fMRI signal return to baseline. For these reasons, the quantification of TTPs and response magnitudes should be seen as tentative.

The BOLD and VASO TTPs are shown in Figure 6A. For the BOLD initial peak, we found longest TTP at closer distances and a trend of decreasing TTPs with distance. For the BOLD trough, we found longest TTPs for the closest distance bin, with similar TTPs for distance bins *>* 8-10 mm. In contrast, the BOLD post-stimulus peak showed shortest TTP for the closest distance bin (6-8 mm) and increasing TTPs for more distant bins. BOLD TTPs across layers were mainly stable within individual distance bins, only showing a systematic difference for the post-stimulus peak. Specifically, for distances *>* 8-10 mm, the superficial layer peaked slightly earlier than middle and deep layers.

**Figure 6:**
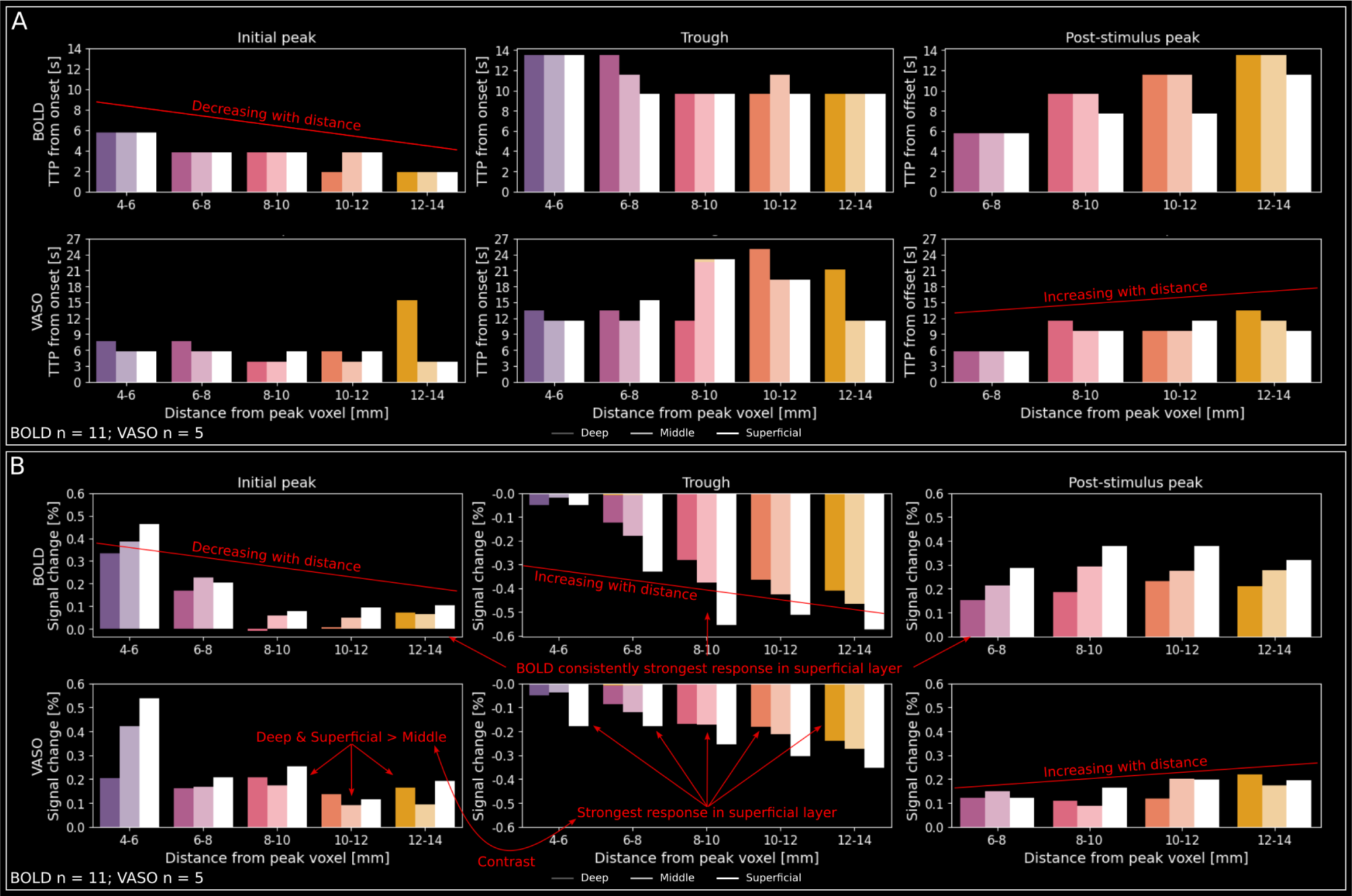
T**e**ntative **quantification of the triphasic response. A** BOLD (top; n = 11) and VASO (bottom; n = 5) times to peak (TTP) of the initial peak (left), trough (middle) and post-stimulus peak (right) of the triphasic response for distance bins *≥* 4-6 mm. Note that for the post-stimulus peak, we only took distances *≥* 6-8 mm into account because for smaller distances, we did not observe a corresponding response in the group results. For the initial peak and trough, times were taken with respect to stimulus onset. For the post-stimulus peak, times were taken from stimulus offset. Plotted is the mean across participants. Due to the high variation across participants, we refrained from plotting error bars and the layer-results did not reach significance and should be interpreted with caution. General trends and noteworthy features are indicated by the annotation. B) BOLD (top; n = 11) and VASO (bottom; n = 5) signal changes (%) of the initial peak (left), trough (middle) and post-stimulus peak (right) of the triphasic response for distance bins *≥* 4-6 mm. General trends and noteworthy features are indicated by the annotation.

For the VASO TTPs, we generally found similar patterns as for BOLD, however with more variability. Noteworthy differences with respect to the BOLD data are as follows: For the VASO initial peak, we found more stable TTPs across distance bins across layers (except for a surprisingly long TTPs for deep layers in the 12-14 mm distance bin, likely due to noise). For the VASO trough, we found increasing TTPs for distance bins between 4-6 and 10-12 mm, followed by again shorter TTPs for the 12-14 mm bin. Across layers, the most systematic difference between BOLD and VASO was slightly longer VASO TTPs for deep layers in 10 out of 14 layer comparisons, which we never found for BOLD.

The respective BOLD and VASO signal changes are shown in Figure 6B. For the BOLD initial peak, we found highest signal changes at closer distances and decreasing peak strengths for increasing distance until the 8-10 mm distance bin. For the BOLD trough, we found increasing magnitudes (more negative signal changes) for distances between 4-6 and 8-10 mm, with only minor variation in the responses for distance bins *>* 8-10 mm. For the BOLD poststimulus peak, we found a trend towards increasing signal changes with distance, except for the most distant bin. No clear layer-effect could be seen with BOLD. Rather, strongest activation was generally found in superficial layers, as expected based on the draining vein effect (with only two minor exceptions that will not be discussed further).

For the VASO signal changes, we found similar patterns over distance bins as for BOLD. Generally, the trends of decreasing initial peak magnitude, increasing trough magnitude (more negative) and increasing post-stimulus peak magnitude remained. In contrast to BOLD, we found some more consistent layer-differences across distance bins. Specifically, for the initial VASO peak across layers, we found systematically higher signals in superficial and deep layers compared to middle layers for distance bins *>* 8-10 mm. In contrast, we consistently found highest VASO trough magnitudes in superficial layers, irrespective of distance. For the poststimulus peak, we did not find any consistent pattern across layers between distance bins.

## Discussion

### Summary of results

In the present study, we explored the laminar and spatio-temporal BOLD and CBV response-characteristics to passive vibrotactile stimulation (30 seconds duration) in the human primary somatosensory cortex. To do so, we acquired depth-resolved (0.75 mm in-plane resolution) BOLD and CBV data using VASO at 7T (f)MRI. For BOLD, we found reliable digit representations in putative BA3b for all participants. For VASO, we found identifiable representations for all 3 digits in 7 out of 11 participants, and for 2 out of 3 digits in 2 further participants. Crucially, in some participants, this digit mapping using VASO was possible with limited data of as few as 4 repetitions per digit. Furthermore, we found that the responses to bottom-up digit stimulation across cortical depth followed the expected pattern of highest activation between middle and superficial cortical layers for VASO and increasing activation towards the cortical surface for BOLD. Finally, for the first time in humans, we observed a triphasic response to the stimulation of non-preferred digits in the digit-ROIs. Specifically, we found a small initial positive activation, followed by a more sustained negative signal change compared to baseline and a positive post-stimulus response. This pattern was most prominent in the BOLD and to a certain extent also visible in the VASO data. Furthermore, we explored the triphasic pattern as a function of distance from each digit’s peak-activation. We found that the initial peak decreased with distance from the activation hotspot of each digit, and the following deactivation increased with distance from the activation hotspot of each digit. Finally, while tentative, we observed slightly higher VASO signal changes in superficial and deep layers compared to middle layers at higher distances for the initial peak, potentially indicative of feedback-processes.

### Mapping of digit representations using BOLD and VASO

One of the goals of the present study was to investigate to what extent the combined BOLD and VASO acquisition is suitable for obtaining maps of digit-representations in putative BA3b (i.e. the first area to receive detailed sensory input from individual digits; Penfield and Boldrey, 1937). As expected, we found reliable digit-representations based on the BOLD data in all participants (Figures 2 and 3; Supplementary Figures S3-S5) and the overall organization is in agreement with previous studies (Nelson and Chen, 2008; Sanchez-Panchuelo et al., 2010; Sanchez-Panchuelo et al., 2012; Schweizer et al., 2008; Steinbach et al., 2022; Stringer et al., 2011). For the VASO data, we found weaker activation compared to BOLD, as expected due to the challenging SNR (Huber, 2014). Nevertheless, we were able to show activation clusters that matched the location of BOLD activity for all digits in 7 out of 11 participants (Figure 2; Supplementary Figures S3-S5). For 2 additional participants, we found clusters for 2 out of 3 digits that matched the BOLD activation. For 2 out of 11 participants we found identifiable VASO responses for one digit only. These results seem comparable to those in the only other published study showing VASO responses in multiple individual digit regions of the human somatosensory cortex (de Oliveira et al., 2023). Specifically, the authors showed VASO maps that follow the expected organization in 4 out of 6 participants (*∼* 66 %), which is similar to the *∼*64 % reported here. Future studies will show whether this is due to individual participants’ VASO response characteristics in the somatosensory domain and/or amount of data acquired per participant.

We would like to highlight that the amount of data used in the present experiment is exceptionally small and the VASO results should be interpreted with caution. While we acquired only between 4 and 12 repetitions per digit across participants, for example, Huber et al. (2021b) used 20 blocks of stimulation (30 s, each) with a similar setup. This is almost twice as much as for the participants that underwent the most scanning in our experiment. Furthermore, de Oliveira et al. (2023) acquired a similar amount of data for all participants, as we did for the participants with the most repetitions (approximately 12 repetitions of 30 s per digit). However, de Oliveira et al. (2023) used an active finger flexion task, while we used passive vibrotactile stimulation. Previous studies have shown that the location of digit representations resulting from active and passive tasks is similar, with activation strength being higher for active tasks

(Sanders et al., 2023). Nevertheless, we believe that using passive vibrotactile stimulation is crucial for more applied experimental settings because it allows consistently accurate, experimenter controlled administration of stimuli, compared to more variable active tasks or manual brushing. Finally, despite the small amount of data, a key benefit of the VASO-sequence is that both BOLD- and CBV-weighted images are acquired in an interleaved fashion. As a result, the high sensitivity of BOLD can be used to guide the definition of ROIs, to then extract additional depth-resolved CBV responses with higher specificity.

With the restrictions of the limited VASO data in mind, we still extend the literature by providing insights into laminar spatial and temporal dynamics in response to passive stimulation of individual digits in humans. When investigating responses to stimulation of the preferred digit within the digit ROIs across cortical depth, unsurprisingly, we found the BOLD response to increase towards the cortical surface due to the well known draining-vein effect (Figure 3B; De Martino et al., 2013; Polimeni et al., 2010; Turner, 2002). On the other hand, the VASO response showed a peak between middle and superficial cortical depths (Figure 3B). At first glance, the VASO peak response is located surprisingly close to the cortical surface compared to an expected response centered in gray matter as reported in VASO studies in the visual domain (e.g. Akbari et al., 2022; Dresbach et al., 2023). In part, this may be explained by the low cortical thickness of the primary somatosensory cortex (Fischl and Dale, 2000). Furthermore, because we found clear activation for all digits of all participants in BOLD but not VASO, we based our ROIs on the BOLD activation, which may include a larger amount of surface vessels that will bias the VASO response towards CSF. Finally, we did not employ an attention task in the present study. Therefore, it is conceivable that participants actively directed their attention towards the stimulation, which has been found to increase responses in superficial layers (e.g. Gau et al., 2020; Lawrence et al., 2019). Future studies in which the participants’ attention is controlled may investigate the laminar CBV response peak.

### Triphasic response as a function of distance

In an exploratory analysis, we found a triphasic BOLD response to stimulation of non-preferred digits within the ROIs (Figure 4). Specifically, we could see a small initial positive response, followed by a sustained signal decrease and a positive response after stimulus cessation. Furthermore, the signal decrease seemed to be larger for more distant digits. Crucially, a signal decrease in response to stimulation of non-preferred digits was barely visible in our results based on the GLM analysis (Figure 3), which highlights the importance of also investigating (laminar) fMRI timecourses in a model-free approach. To investigate this further, we plotted BOLD and VASO signal changes across cortical depth and as a function of distance from the location of strongest activation for each stimulated digit. Generally, the BOLD and VASO magnitudes of the initial peak decreased, while the following trough magnitude increased with distance from the activation center (Figure 5). At distances *≥* 6-8 mm, positive signal changes started to emerge after stimulus cessation.

Generally, similar findings have been reported in regions of the somatosensory cortex representing the rat forepaw (e.g. Devor et al., 2005, 2007), rat whisker (Boorman et al., 2010) and squirrel monkey digits (e.g. Simons et al., 2007) using various invasive methods (e.g. laminar electrode array recordings, voltage sensitive dye imaging, or two photon microscopy). In humans, similar responses have been observed after stimulation of the ipsilateral digits or median nerve using BOLD (e.g. Hlushchuk and Hari, 2006; Kastrup et al., 2008; Klingner et al., 2010, 2011), and/or cerebral blood flow (CBF) measurements (Mullinger et al., 2014; Schäfer et al., 2012). To our knowledge, however, the present study is the first demonstration of a similar contralateral BOLD and CBV response that changes as a function of distance from a positive activation peak, comparable to the animal literature.

Similar responses have been discussed in the literature as potentially resulting from neural and/or vascular mechanisms. With our data, we cannot conclusively differentiate between the two options, which are also not mutually exclusive. However, some aspects of our results may hint more towards a neuronal origin, while other aspects may hint more towards a vascular origin. Therefore, we will discuss both options with respect to the initial peak, trough, and post-stimulus peak.

### Initial peak

A small initial positive BOLD response in regions with an overall negative response have been reported in both animals (e.g. Devor et al., 2007; Shmuel et al., 2006) and humans (e.g. Kastrup et al., 2008; Klingner et al., 2011), albeit not unanimously (for example, see Hlushchuk and Hari, 2006). Klingner et al. (2011) argue that this initial peak may be due to a decrease in the cerebral metabolic rate of oxygen (CMRO_2_) with delayed reductions of CBF that lead to the negative BOLD response. This would match the findings by Shmuel et al. (2006), who see a small BOLD increase before a larger negative response in the absence of increased neural activity during the initial increase. However, a decrease in CMRO_2_ alone is unlikely to explain our data because we see a corresponding CBV increase in our VASO data. As discussed above, interpretations of the VASO data in the present study have to be treated with caution due to the challenging noise level and high variability across participants. However, the initial VASO peak is the most consistent feature we observe across distance bins. Therefore, our data could hint towards a vascular origin of the initial positive response.

For example, it is conceivable that the same artery feeds the active digit and neighboring (later inhibited) cortical regions, which may lead to a temporary signal increase along the feeding pathway. This would be supported by the slightly shorter times to peak of the initial BOLD and VASO responses of the more distant, compared to closer bins (Figure 6A). In future studies, this could be tested by using vascular information to delineate arteries and investigate whether the initial peak of the triphasic response is stronger in the proximity of an artery feeding the digit region. However, next to vascular effects, our data could also hint towards a neural origin of the initial positive response. Specifically, we found the small initial VASO response to be slightly higher in superficial and deep, compared to middle layers for more distant bins (*≥* 8-10 mm; Figure 6B). Commonly, it is assumed that superficial and deep layers receive feedback information from other cortical regions (Felleman and Van Essen, 1991). Potentially, this feedback input could then in turn trigger the following inhibitory response. This possibility would match the findings by Boorman et al. (2010), who report an initial short-latency electrophysiological response to stimulation in the area surrounding the main positive response. However, this would beg the question of why we found the earlier times to peak of the initial response in more distant regions than closer to the site of activation. While this could be due to simultaneous input from the thalamus, these interpretations are highly speculative, especially given that the laminar differences are very small and do not survive statistical testing.

### Trough

After the small initial signal increase, we observed sustained negative BOLD and VASO signal changes that showed a trend towards larger magnitudes with increasing distance from the activation center. This finding is in line with several animal studies (e.g. Devor et al., 2005, 2007).

Early studies suggested that negative fMRI responses originated from CBF redirection to active areas, leaving non-active areas with lower CBF and therefore negative BOLD signal changes (“blood steal”, Harel et al., 2002; Woolsey et al., 1996). In our data, the negative BOLD and VASO signal changes showed a monotonic increase from deep to superficial layers, in accordance with laminar differences in baseline blood volume (Boorman et al., 2010; Huber et al., 2014; Huber et al., 2021b). Therefore, it is possible that the negative BOLD and VASO signal changes have a vascular origin. However, more recent literature strongly indicates that blood-stealing is unlikely to solely explain negative fMRI signal changes (Boorman et al., 2010; Huber et al., 2014; Kennerley et al., 2012; Pasley et al., 2007; Shmuel et al., 2006; Stefanovic et al., 2004). Therefore, we propose that our results could also be explained by neuronal inhibition.

For example, Kastrup et al. (2008) showed that participants’ sensitivity to a stimulus was reduced in regions that also showed a negative BOLD response. Specifically, the authors found that the magnitude of the negative BOLD response in the ipsilateral somatosensory cortex (evoked by stimulation of the right median nerve) correlated with increases in a current perception threshold when simultaneously stimulating the contralateral (left) index finger (Kastrup et al., 2008). In other words, participants with a more negative BOLD response required higher currents to perceive a second stimulus, indicative of functional inhibition. Without a direct comparison, it is difficult to translate the functional significance of negative ipsilateral responses (as in Kastrup et al., 2008) to the negative responses in the contralateral surround reported here. However, Boorman et al., 2010 investigated the center-surround responses in the contraas well as the ipsilateral hemisphere of the rat somatosensory cortex and found that the negative BOLD signal changes in the contralateral surround resembled those in the ipsilateral hemisphere. Given that the haemodynamic processes in ipsi- and contralateral hemispheres seem to be similar, we hypothesize that the negative BOLD and CBV responses reported here, may also reflect similar inhibitory neuronal processes as reported in Kastrup et al. (2008).

Another study that may hint towards a neural origin of the negative haemodynamic responses reported here was performed by Simons et al. (2007), in which the authors used optical imaging spectroscopy in squirrel monkeys and showed surround suppression for vibrotactile stimuli. Interestingly, the site of largest negativity was not uniform, but overlapped with regions that would usually be co-stimulated in the natural environment. This could mean that the functional architecture and not the vasculature is determinant for the negative signal changes. On the one hand, it is also conceivable that regions that are more often co-activated are supplied by the same vascular structures, which would make the cerebral support system more efficient. Still, assuming that different digits are more often co-activated with other digits than with the face (represented laterally of the digits in the primary somatosensory cortex), this leads to the interesting hypothesis that the inhibition during stimulation of D3 should be more uniform across directions than for D1 or D5. Future studies could investigate this further by combining vascular information with behavioral tasks as in Kastrup et al. (2008).

### Post-stimulus peak

After the stimulus cessation, we found an overshoot following negative BOLD and VASO responses in more distant bins (white arrows in Figure 5B). This finding is comparable to results in the human visual cortex as measured with CBV, BOLD and CBF (Huber et al., 2014), human ipsilateral somatosensory cortex as measured with BOLD (Kastrup et al., 2008) and the rat barrel cortex as measured with BOLD, CBF (as measured with laser Doppler flowmetry), and total CBV (as measured with 2D-optical imaging spectroscopy; Boorman et al., 2010).

In our results, the timing of the post stimulus response as a function of distance from the activation center may hint towards a vascular origin. Specifically, while we found decreasing times to peak with distance from the activation center for the initial peak, we found increasing times to peak with distance from the activation center for the post-stimulus peak (Figure 6A). Before, we argued that the decreasing times to peak with distance from the activation center for the initial peak could be indicative of common supply routes for different digit representations. Likewise, a similar draining architecture could be in place, thus leading to the opposite pattern at stimulus cessation.

On the other hand, Shmuel et al. (2006) report a similar post-stimulus BOLD peak in regions with prior negative BOLD signal changes in the monkey visual cortex. Crucially, the simultaneously recorded neural activity convolved with a haemodynamic response function is in close agreement with the BOLD response. Therefore, while we cannot rule out vascular influences, a neural origin of the post-stimulus seems likely. However, the functional significance of the post-stimulus increase is still to be elucidated.

## Limitations and future directions

There are several limitations that have implications for the results discussed in the present manuscript.

Most importantly, the data presented were collected in the context of a larger study. Therefore, we were only able to acquire a limited amount of stimulation data for some participants (4 stimulation periods of 30 s per digit). This is especially detrimental for the VASO data, given its challenging SNR and low temporal efficiency. Specifically, other studies with a similar setup (e.g. Huber et al., 2021b) acquired almost twice as much data (20 stimulation periods of 30 s) compared to the participants for which we acquired the most data (12 stimulation periods of 30 s per digit). Nonetheless, we were able to obtain reasonable responses to digit stimulation in most participants, even in the VASO data. However, there is considerable variability in the data across participants, regarding the triphasic response. Therefore, the results in this study have to be interpreted with caution and future studies will have to validate the present results with appropriate statistical testing.

Furthermore, the inter-trial period of our design did not allow a complete return to baseline for the BOLD and VASO responses. We employed a 30 s on/off block design, which is used in many layer-fMRI studies, especially when also recording non-BOLD responses, because of a good tradeoff between detection sensitivity, temporal efficiency and signal integrity (e.g. Dresbach et al., 2023; Faes et al., 2023; Huber et al., 2017; Pizzuti et al., 2023). Given that the triphasic response observed here is clearly locked to the stimulus onset, we do not believe that the effect is driven by preceding stimuli. However, because of partially overlapping responses, the exact quantification of the response timings and magnitudes in the event-related averages is difficult. Also, the baseline calculation in the event-related averages may have slightly overestimated the baseline value. Therefore, signal changes of the peaks may be slightly underestimated, while the signal changes of the troughs may be slightly overestimated. In future studies, a longer inter-stimulus period could be used to shed further light on the timing and magnitudes of the observed responses.

Next to improving on the points discussed above, future studies could acquire dedicated vascular imaging. This could help to examine whether different aspects of the triphasic response reported here are more likely to have a neural or a vascular origin. In animal studies reporting this phenomenon (Devor et al., 2007, 2008), invasive methods are available to investigate neuronal spiking and local field potentials alongside haemodynamic signals and arterial spiking. While these methodologies are largely unavailable in humans, for example, multi-echo gradient-echo images could be acquired at ultra-high resolutions to delineate individual arteries and veins and then plot time resolved BOLD and VASO activation (Dresbach et al., 2024; Gulban et al., 2022).

Furthermore, the functional data coverage could be expanded to include the ipsilateral somatosensory cortex. This way, the triphasic contralateral responses observed in this study could be compared to the human literature that has shown similar effects following ipsilateral stimulation.

Finally, while the neurovascular mechanisms of the different aspects of the triphasic response have been well described in the animal literature, their functional significance is still relatively unclear. Including behavioral testing (similar to Kastrup et al., 2008) in humans, could shed light on the functional significance.

## Conclusion

We characterized haemodynamic responses to stimulation of individual digit-tips across cortical depth at 0.75 mm in plane spatial resolution using BOLD and VASO fMRI. We could identify digit-specific ROIs in putative Brodmann area 3b, following the known anatomical organization. In the ROIs, the BOLD response increased towards the cortical surface due to the draining vein effect, while the VASO response was more shifted towards middle cortical layers, likely reflecting bottom-up input from the thalamus, as expected. Finally, we explored the temporal signal dynamics for BOLD and VASO as a function of distance from activation peaks resulting from stimulation of contralateral digits. With this analysis, we observed a triphasic response consisting of an initial peak and a subsequent negative deflection during stimulation, followed by a positive post-stimulus response in BOLD and to some extent in VASO. Interestingly, the initial peak was most consistently found for VASO and seemed to have a more feed-back layer signature for surrounding areas. While similar responses were reported with invasive methods in animal models, to our knowledge, this is the first demonstration of potential neuronal excitation-inhibition in a center-surround architecture across layers in the human somatosensory cortex. Given that, unlike in animals, human experiments do not rely on anesthesia and can readily implement extensive behavioral testing, obtaining this effect in humans is an important step towards further uncovering the functional significance of the different aspects of the triphasic response.

## Acknowledgments

We thank Stefano Moia for discussions about potential explanations for the triphasic response, Vojťech Smekal for helpful discussions on the manuscript and Renate Schweizer for discussions regarding finger mapping, analysis and interpretation. Special thanks goes to the remaining ”Maastricht layer-seminar” members Ana Arsenovic, Lonike Faes, Lasse Knudsen and Kenshu Koiso, Jan Kurzawski, Alessandra Pizzuti, Yawen Wang for countless discussions on topics related to layer-fMRI.

SD is supported by the ‘Robin Hood’ fund of the Faculty of Psychology and Neuroscience and the department of Cognitive Neuroscience. RG is partly funded by the European Research Council Grant ERC-2010-AdG269853 and Human Brain Project Grant FP7-ICT-2013-FET-F/604102. OFG and JE are funded by Brain Innovation. NW has received funding from the European Research Council under the European Union’s Seventh Framework Programme (FP7/2007-2013) / ERC grant agreement n° 616905; from the European Union’s Horizon 2020 research and innovation programme under the grant agreement No 681094; from the Deutsche Forschungsgemeinschaft (DFG, German Research Foundation) – project no. 347592254 (WE 5046/4-2); from the Federal Ministry of Education and Research (BMBF) under support code 01ED2210. Renzo Huber was supported by the NIMH Intramural Research Program (#ZI- AMH002783) and by NWO VENI project 016.Veni.198.032 during this project.

## Data and Software availability statement

Analysis code is available on GitHub: *<*https://github.com/sdres/s1Anfunco*>*. Raw data is available on OpenNeuro: *<*https://openneuro.org/datasets/ds005238/versions/1.0.0*>*.

## Author Contributions

|  |  |
| --- | --- |
| Conceptualization | SD, TS, JE, AK, RG, LH |
| Data curation | SD, TS, JE, AK, LH |
| Formal analysis | SD, OFG, SK |
| Funding acquisition | SD, AK, RG, LH |
| Investigation | SD, OFG, TS, JE, LH |
| Methodology | SD, OFG, TS, NW, RG, LH |
| Project administration | SD, RG, LH |
| Resources | SD, AK, RG, LH |
| Software | SD, OFG, SK, RG, LH. |
| Supervision | OFG, AK, NW, RH, LH |
| Validation | SD |
| Visualization | SD, OFG, RG, LH |
| Writing - original draft | SD |
| Writing - review and editing | SD, OFG, TS, JE, SK, NW, RG, LH |

## Supplementary Materials

**Figure S1:**
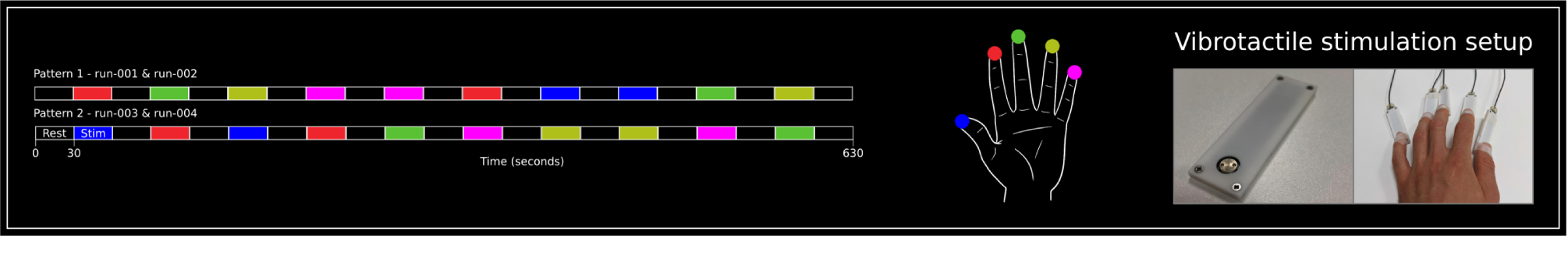
Stimulation setup of participant sub-12. For this participant, we stimulated the thumb (D1) and pinky (D5) of the left hand, in addition to D2-D4. Increasing the duration of each run to accommodate the stimulation of additional digits would have been detrimental for participant comfort. Therefore, we reduced the run duration by only including 2 repetitions/digit in each run (resulting in a run duration of 10.5 minutes, including 30 seconds of initial rest). To account for the reduction of repetitions per digit, we acquired 4 runs of stimulation for this participant, resulting in 8 repetitions/digit. Also here, we generated 2 stimulation patterns. For run-001 and run-002 we used pattern 1, while for run-003 and run-004 we used pattern 2.

**Figure S2:**
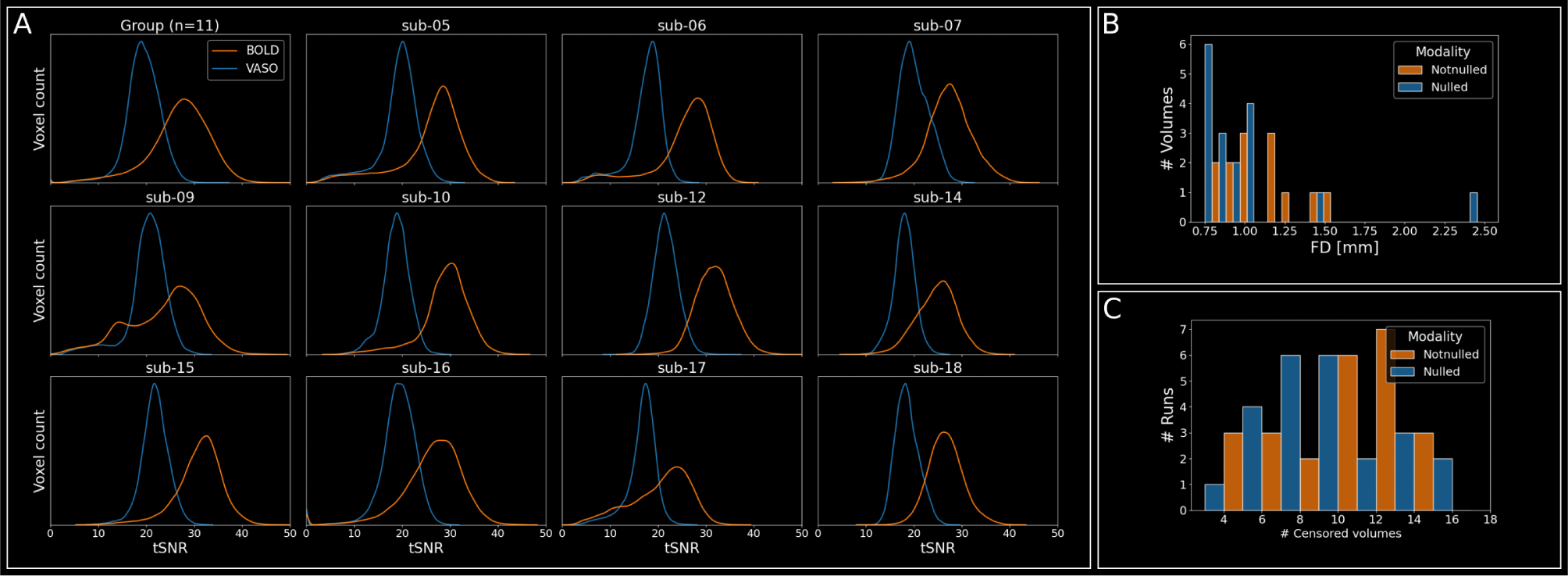
Quality assessment A. TSNR values for BOLD and VASO on the group level (n = 11) and for each participant separately. Values were extracted from the “perimeter chunk” generated by the *LN2 MULTILATERATE* program (i.e. the geodesic disc propagated across cortical depth from in which digit specific ROIs were generated; see Figure 3A). **B** Histogram of the number of volumes showing framewise displacement (FD) *>* 0.75 mm (in-plane voxel resolution) of all participants and runs for nulled and notnulled data separately. A total of 9198 volumes (nulled and notnulled combined) was acquired across participants. Of those, 30 volumes (¡0.33%) showed (FDs) greater than our voxel size (0.75 mm). 13 (in 6 individual runs) of those were in notnulled and 17 (in 9 individual runs) were in nulled time series. **C** Histogram of the number of runs with a certain number of censored volumes for nulled and notnulled timeseries separately as given by the *fsl motion outliers* program.

**Figure S3:**
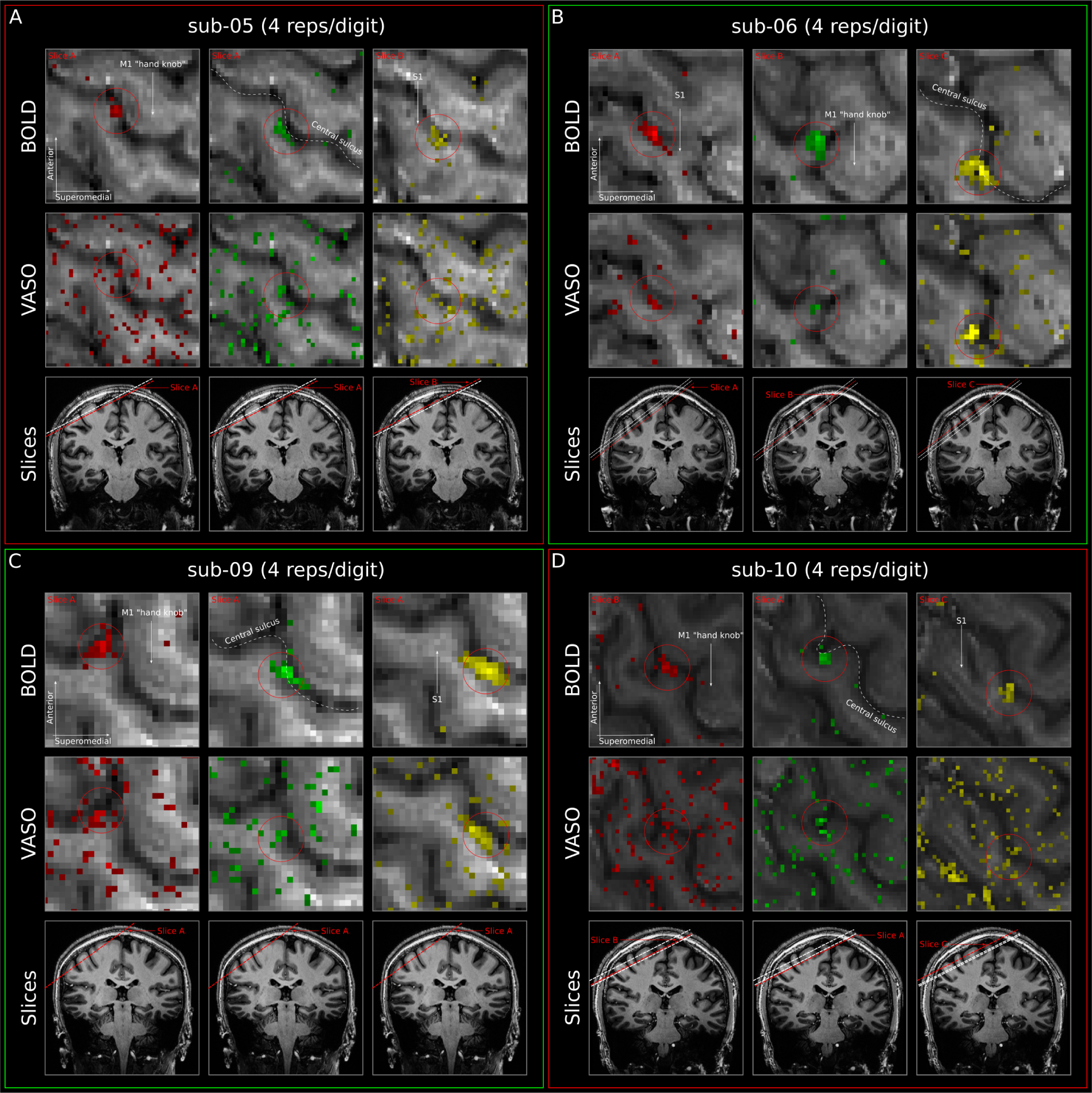
Stimulation results for participants sub-05-sub-10. Same as Figure 2B but for participants sub-05 (**A**), sub-06 (**B**), sub-09 (**C**), and sub-10 (**D**). A green box around the participant’s plots indicates that we could identify 3/3 digit representations based on the VASO data, an orange box around the participant’s plots indicates that we could identify 2/3 digit representations based on the VASO data, and a red box around the participant’s plots indicates that we could identify only 1/3 digit representations based on the VASO data. Colorbars and identification of digit-clusters are the same as in 2

**Figure S4:**
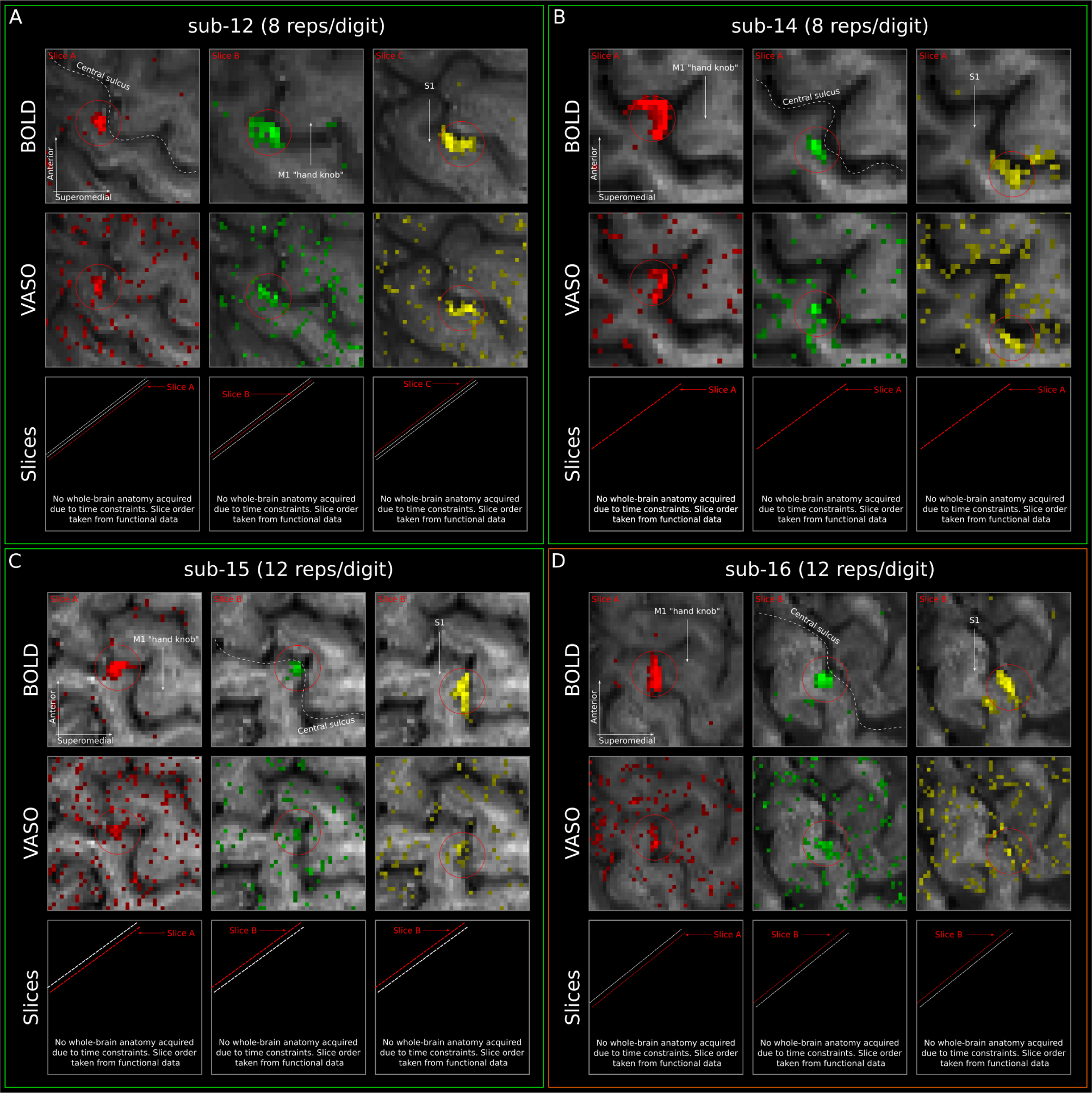
Stimulation results for participants sub-12-sub-16. Same as Figure 2B but for participants sub-12 (**A**), sub-14 (**B**), sub-15 (**C**), and sub-16 (**D**). A green box around the participant’s plots indicates that we could identify 3/3 digit representations based on the VASO data, an orange box around the participant’s plots indicates that we could identify 2/3 digit representations based on the VASO data, and a red box around the participant’s plots indicates that we could identify only 1/3 digit representations based on the VASO data. Colorbars and identification of digit-clusters are the same as in 2

**Figure S5:**
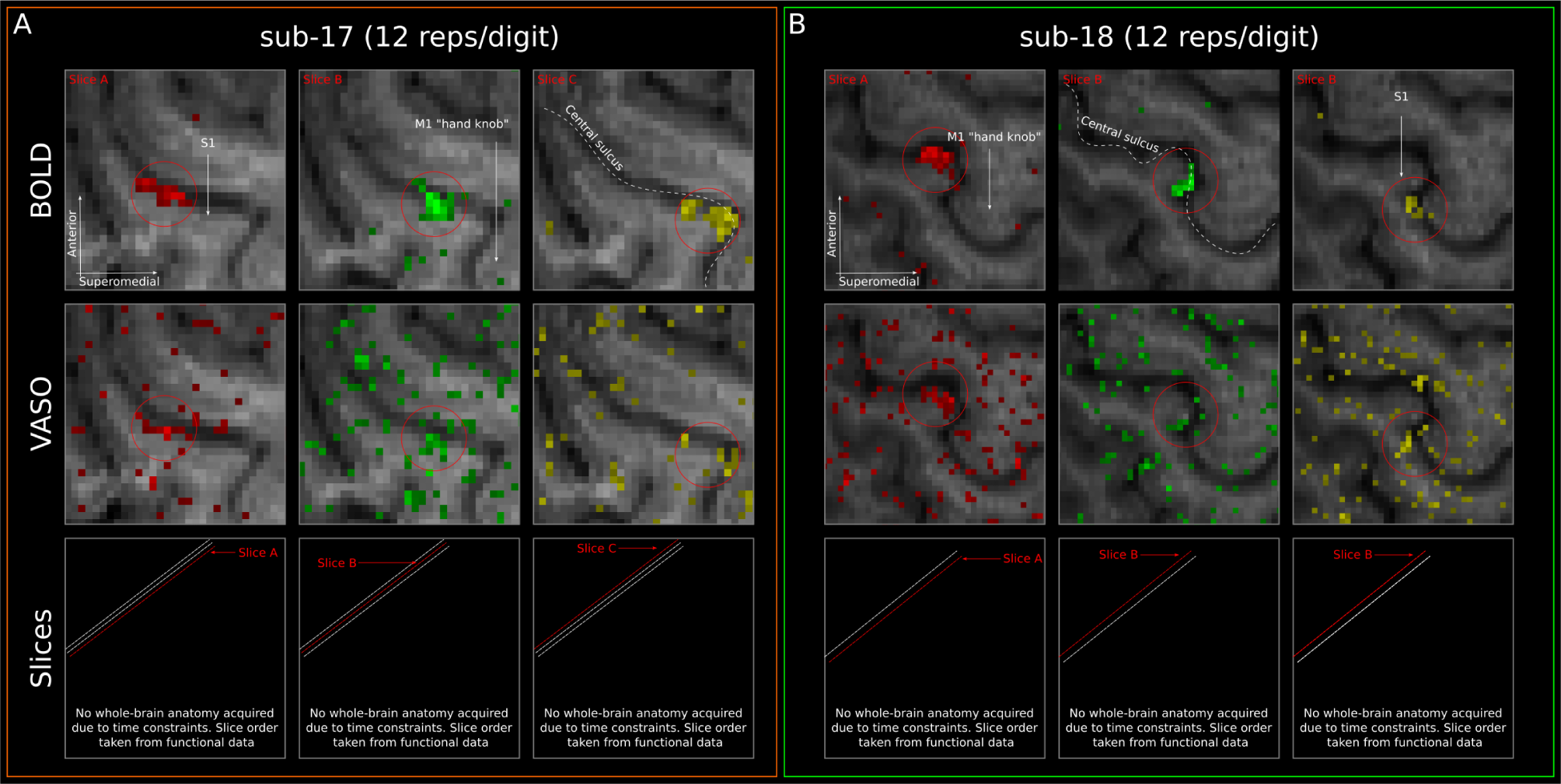
Stimulation results for participants sub-17 & sub-18. Same as Figure 2B but for participants sub-17 (**A**) and sub-18 (**B**). A green box around the participant’s plots indicates that we could identify 3/3 digit representations based on the VASO data, an orange box around the participant’s plots indicates that we could identify 2/3 digit representations based on the VASO data, and a red box around the participant’s plots indicates that we could identify only 1/3 digit representations based on the VASO data. Colorbars and identification of digit-clusters are the same as in 2

**Figure S6:**
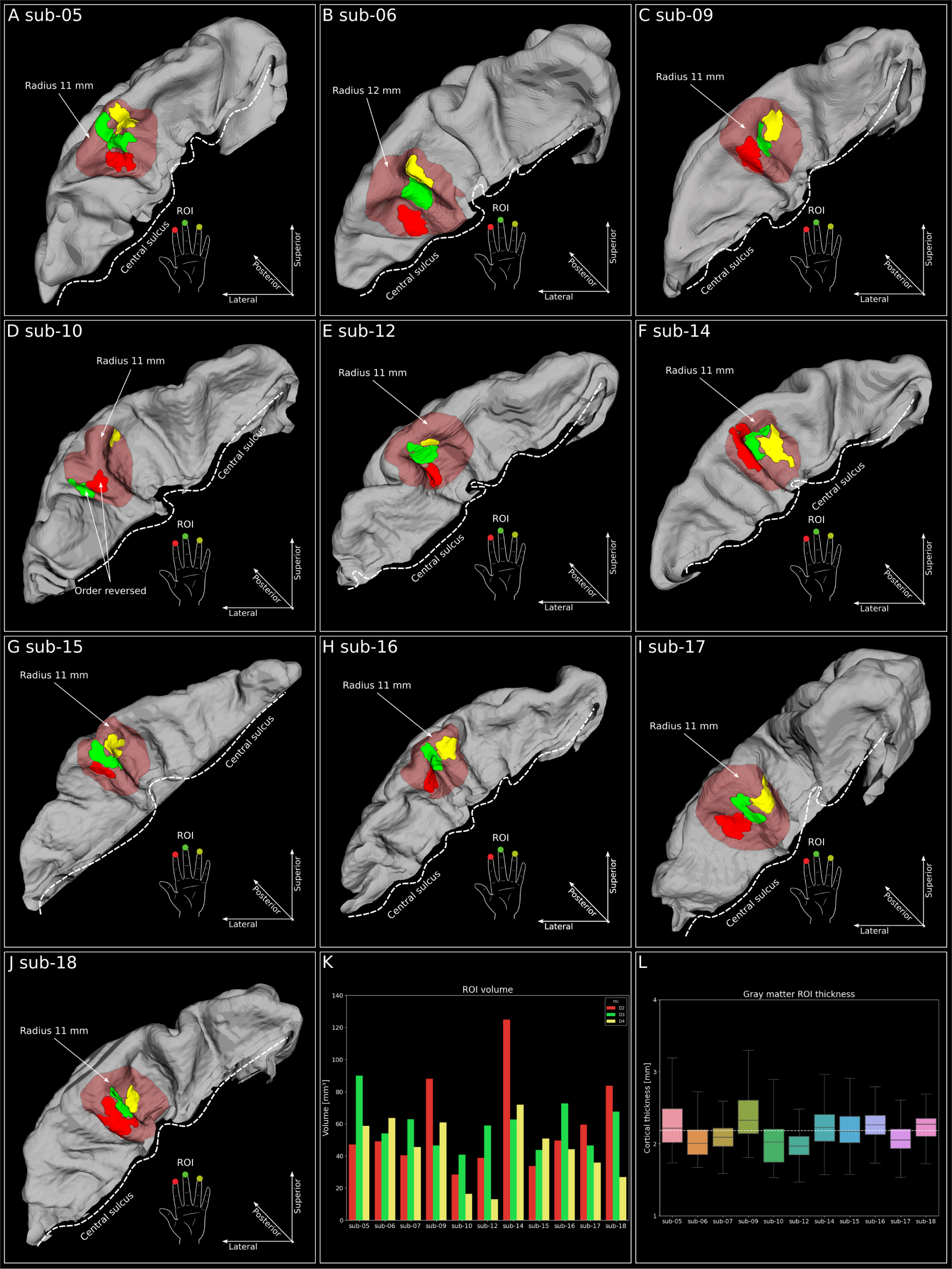
ROIs of individual participants. **A-J**) ROIs of individual participants. **K** ROI volume (mm3) of individual participants for the 3 digits separately. **L** Voxel-wise cortical thickness collapsed over ROIs for all participants independently. Dashed white line indicates mean across participants (*∼*2.1 mm). The box shows the quartiles, whereas whiskers show the rest of the distribution across all voxels.

**Figure S7:**
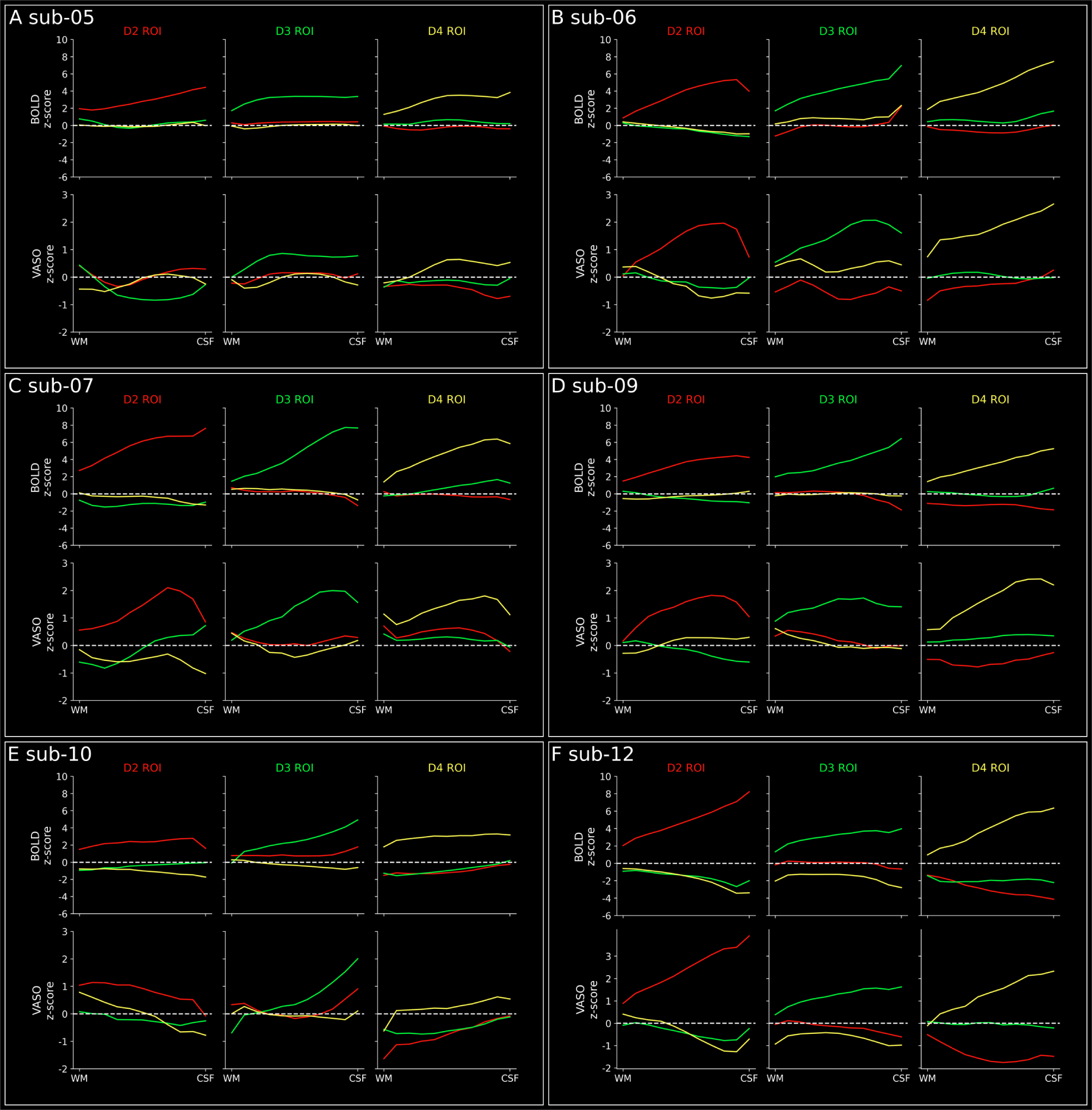
**BOLD and VASO activation across cortical within ROIs based on BOLD for participants sub-05 - sub-12.**

**Figure S8:**
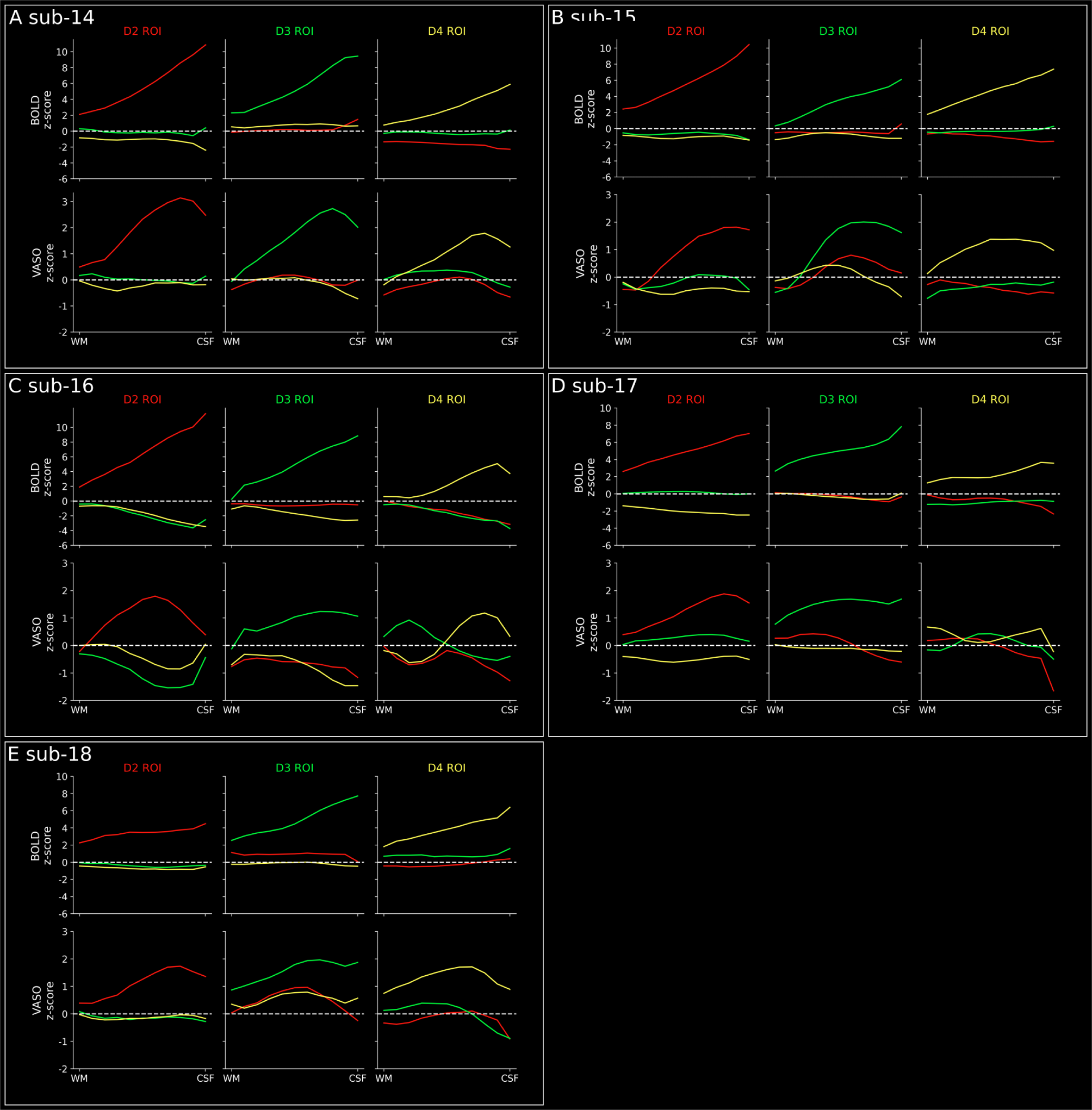
**BOLD and VASO activation across cortical within ROIs based on BOLD for participants sub-14 - sub-18.**

**Figure S9:**
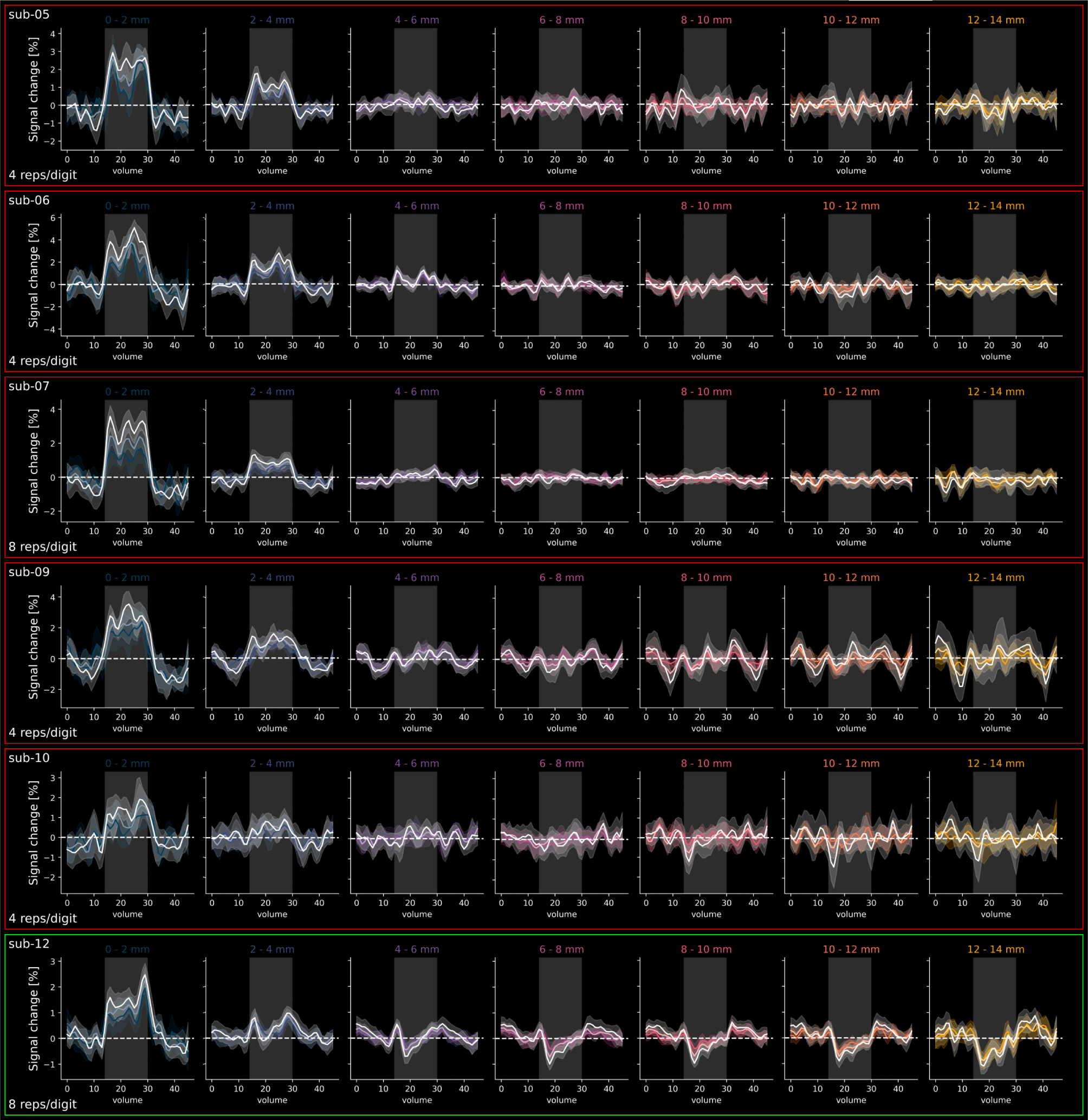
BOLD results of distance analysis for individual participants. Same as Figure 5B, but for participants sub-05, sub-06, sub-07, sub-09, sub-10 & sub-12 individually. A red box around a participant’s plots indicates that we did not find indications of a triphasic response for this participant. A green box around a participant’s plots indicates that we did find indications of a triphasic response for this participant.

**Figure S10:**
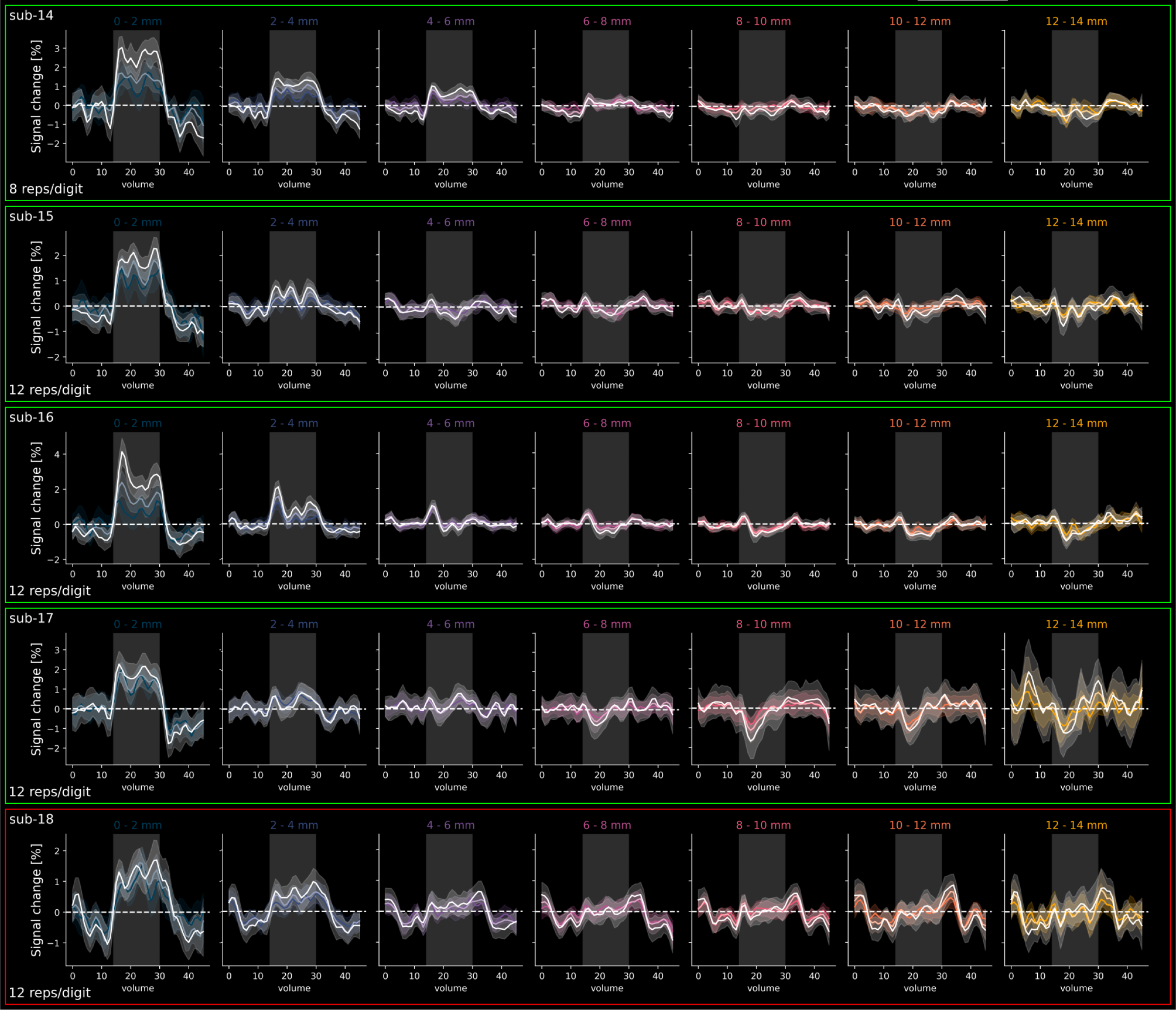
BOLD results of distance analysis for individual participants. Same as Figure 5B, but for participants sub-14, sub-15, sub-16, sub-17, and sub-18 individually. A red box around a participant’s plots indicates that we did not find indications of a triphasic response for this participant. A green box around a participant’s plots indicates that we did find indications of a triphasic response for this participant.

**Figure S11:**
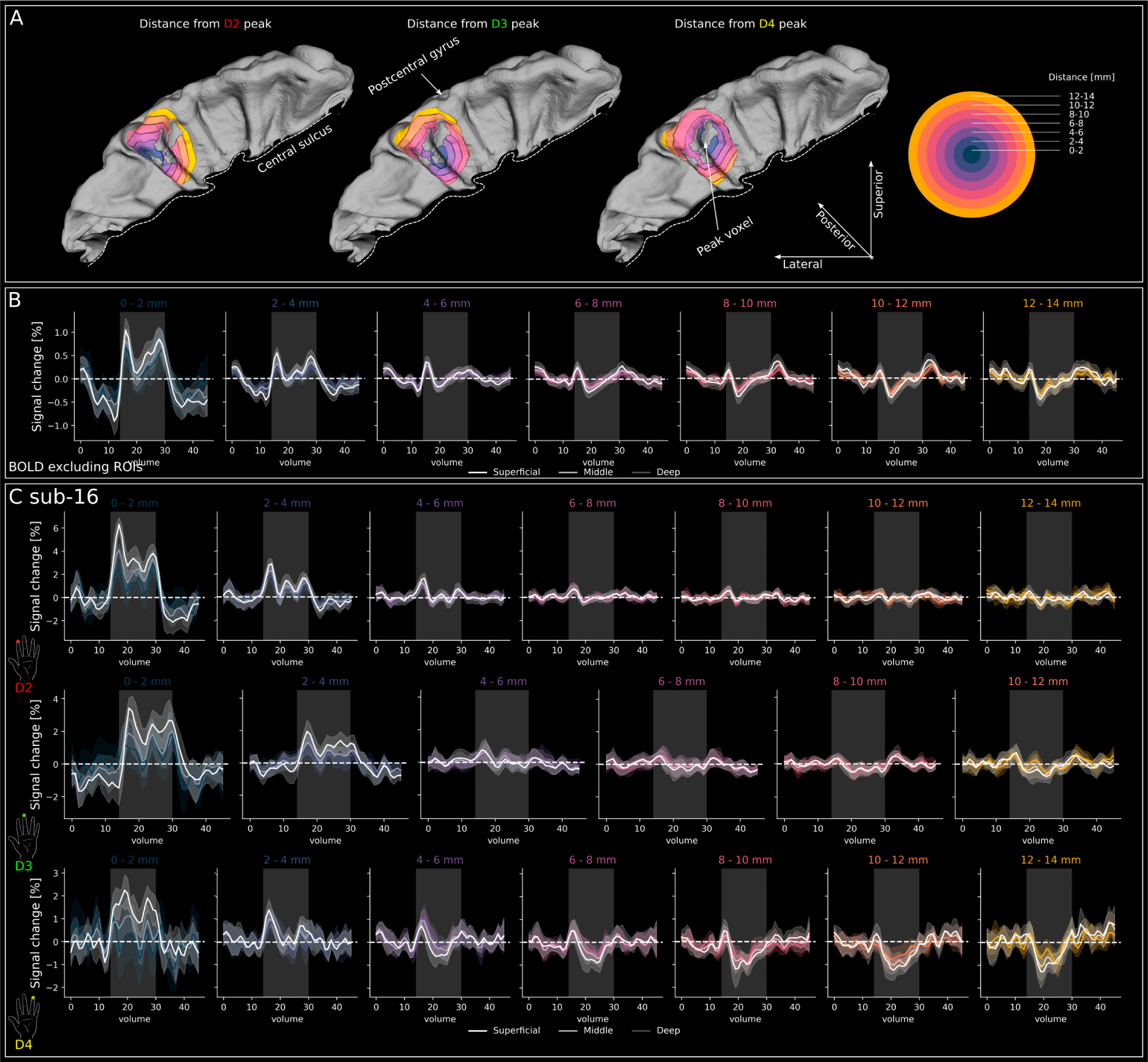
Additional distance-dependent analyses A. Same as Figure 5A but showing the distance bins excluding the digit ROIs for participant sub-07. **B** Same as Figure 5B but excluding signal from digit ROIs. Crucially, the triphasic response is preserved. Note that the scale on the y-axis is changed with respect to Figure 5B, which makes the positive and negative deflections appear to be larger for distance bins *>* 2 mm. **C** Same as Figure 5A but for distance bins with respect to the 3 digits individually in one participant (sub-16). Crucially, the initial triphasic response is preserved in all individual digits. Note that the scale on the y-axis is changed between digits and, therefore, the magnitude of the positive and negative deflections appear different between rows.

**Figure S12:**
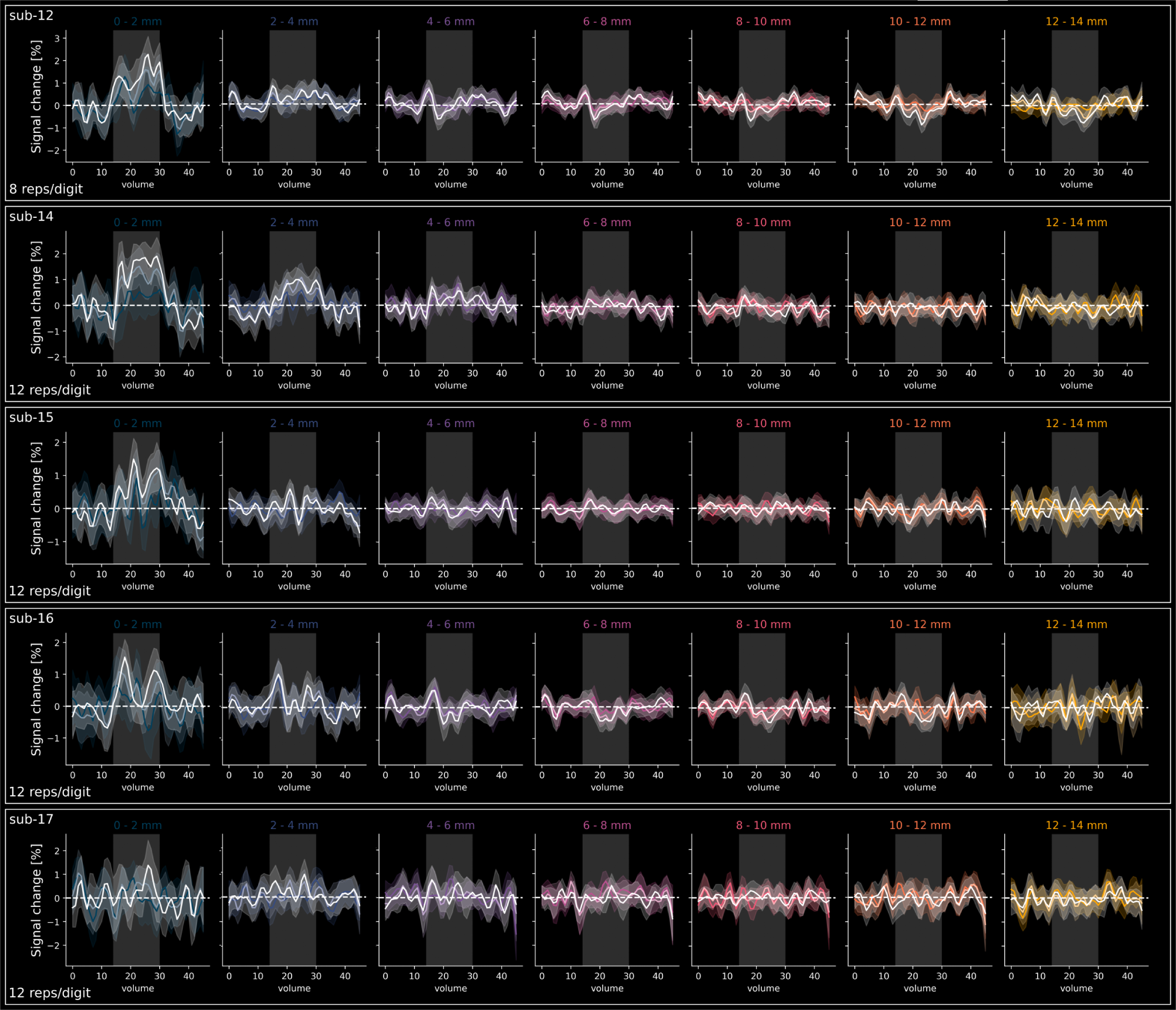
VASO results of distance analysis for individual participants. Same as Figure 5B, but for the subset of individual participants that showed indications of the triphasic response in BOLD (sub-12, sub-14, sub-15, sub-16 & sub-17).

**Table 1:** Session overview of participants.

| Participant | Repetitions/digit | Digits stimulated | MP2RAGE slab images |
| --- | --- | --- | --- |
| sub-01-sub-04 | Pilots | - | - |
| sub-05 | 4 | 3 | 3 |
| sub-06 | 4 | 3 | 3 |
| sub-07 | 8 | 3 | 3 |
| sub-08 | Session aborted due to participant discomfort | - | 3 |
| sub-09 | 4 | 3 | 3 |
| sub-10 | 4 | 3 | 3 |
| sub-11 | Problems during scanning | - | - |
| sub-12 | 8 | 5 | 3 |
| sub-13 | Problems during scanning | - | - |
| sub-14 | 8 | 3 | 3 |
| sub-15 | 12 | 3 | 3 |
| sub-16 | 12 | 3 | 3 |
| sub-17 | 12 | 3 | 3 |
| sub-18 | 12 | 3 | 3 |

